# Deeper neuronal and glial proteomic insights using an optimized pipeline for proximity labeling proteomics

**DOI:** 10.64898/2026.01.21.700735

**Authors:** Wooyoung Eric Jang, Upasna Srivastava, Amanda D. Brandelli, Prateek Kumar, Claudia Espinosa-Garcia, Dilpreet Kour, Rashmi Kumari, Srikant Rangaraju

## Abstract

Proximity-based proteomics using TurboID has enabled cell-type–specific profiling without the need for cell purification, although major bottlenecks in sample lysis, biotinylated protein enrichment, digestion, and mass spectrometry (MS) parameters have limited depth of proteome coverage. Here, we systematically optimized these variables using TurboID-based labeling of BV2 microglia in vitro and brain astrocytes in vivo to define conditions that maximize proteome coverage. In microglia, the optimized protocol using 8 M urea lysis with on-bead S-Trap digestion and data-independent acquisition MS (DIA-MS) identified 4,016 proteins, double the depth of prior studies, and revealed metabolic, ribosomal, lipid-processing, autophagy, and trafficking signatures. Brain astrocyte proteomes were best recovered using SDS lysis with S-Trap digestion and DIA-MS, yielding a proteome of over 3,600 highly enriched proteins, twice the depth of prior astrocyte-TurboID studies. The expanded astrocyte proteomes captured canonical astrocyte markers as well as membrane-associated, vesicular trafficking, and presynaptic protein signatures, consistent with labeling of astrocyte–neuron interface regions, including proteins involved in receptor signaling, lipid metabolism, and plasticity at tripartite synapses, and several AD risk proteins. The increased peptide recovery following S-Trap digestion allowed the reduction of starting material to 20 µg protein for DIA-MS, and enabled multiplexed tandem mass tag (TMT-MS) proteomics using even smaller samples. When applied to synaptosomes enriched from mouse brains with neuronal TurboID labeling, our pipeline identified a synapse-specific proteome of 2,529 proteins, revealing synaptic, mitochondrial and disease-relevant signatures not detectable in prior studies. By tackling critical bottlenecks from tissue processing to MS, our optimized pipelines enable cell-type and compartment-specific proximity-labeling proteomics to obtain comprehensive biological and disease-relevant insights across various biological fields.

## INTRODUCTION

The immense cellular complexity of organs, such as the brain, poses several challenges in understanding the relative contributions of different cell types to homeostatic and disease processes ^1, 2, 3, 4^. While rapid advances in cell type-specific transcriptomics methods allow cellular molecular insights, mRNA-level measurements modestly reflect functionally relevant changes at the protein level ^5, 6, 7^. To address this challenge, proximity labeling techniques using promiscuous biotin ligases to label cellular proteomes of specific cell types in native tissues are being rapidly adopted across different biological contexts and model systems including mice ^8^. The major advantage of these proteome labeling strategies is the ability to obtain cell type-specific proteome insights without requiring purification of individual cell types that use harsh dissociation methods that perturb cell state or cause cellular disruption. TurboID represents a bioengineered biotin ligase that has high catalytic efficiency that enables robust proximity labeling of proteins in specific sub-cellular domains, cellular compartments (via TurboID fusion to localization sequences); or broad cellular proteomic labeling (without localization tags) ^9, 10^. Our group recently adopted the TurboID approach using genetic Cre/lox approaches to label and profile the cytosolic proteomes of excitatory and inhibitory neurons as well as astrocytes ^11, 12^. This approach, called cell type specific biotinylation of proteins (CIBOP) and related proximity labeling methods have broad applicability across tissue types, cell types and sub-cellular compartments in mammalian in vitro and in vivo model systems ^11, 13^.

The CIBOP approach relies on the Cre–Lox system for precise genetic control, which affords spatiotemporal resolution, enabling selective biotinylation of the proteome of a single cell type within its native homeostatic context ^11, 13^. As a result, CIBOP provides a molecular snapshot of cell-type-specific proteomes, offering insights into the proteomic signatures and alterations that shape cellular identity. Despite its potential, the practical application of CIBOP presents several technical challenges that impact the depth of proteome coverage by mass spectrometry (MS). Across studies that have used TurboID or closely-related proximity based biotinylation proteomics approaches for cytosolic labeling, proteome coverage has ranged from 1,000-3,000 labeled proteins ^9, 11, 14^. In contrast, the proteome of the whole cell typically contains over 7,000 proteins using label-free quantitative MS methods^15^. This lower coverage by TurboID labeling can be due to several biological and technical reasons. Since a small fraction of the proteome is labeled by TurboID, large amounts of starting material (>300 µg protein) have been needed to allow enrichment of biotinylated proteins to yield sufficient labeled protein for analysis by MS. This often necessitates using large pieces of tissue or pooling samples across experiments, making it challenging to apply current pipelines to small regions or sub-cellular compartments particularly using in vivo model systems. Furthermore, the enrichment of biotinylated proteins using streptavidin bead-based capture inherently has noise due to non-specific binding as well as binding of endogenously biotinylated carboxylases, all of which contribute to background noise in the proteome ^9, 11^. Additionally, streptavidin-coated magnetic beads used for affinity purification (AP) frequently release abundant contaminant streptavidin peptides that dominate MS spectra and suppress detection of target biotinylated peptides. Therefore, rigorous experimental designs incorporating stringent negative controls are essential for confidently assigning cell-type-specific proteomes.

Given the rapid adoption of TurboID and related proximity-based proteomics labeling methods in mouse models and in vitro systems, the scientific communities that use these methods will benefit from a systematic evaluation of each aspect of proximity-based proteomics workflow, including tissue lysis and processing, biotinylated protein enrichment, protein digestion to peptides and cleanup, liquid chromatography (LC) and MS analysis of digested peptides, and finally, data processing and analysis. Protein solubilization, a critical step in sample preparation, employs chaotropic agents (e.g., 8 M urea), multicomponent buffers (e.g., RIPA), or strong anionic detergents such as sodium dodecyl sulfate (SDS), often combined with mechanical disruption. Recent studies suggest that SDS affords superior solubilization efficiency, particularly for membrane-rich biological material ^16, 17^. Moreover, complete solubilization of the postsynaptic density (PSD) complex often requires the combined use of detergents and chaotropes ^18^. However, although SDS enables robust extraction, it is incompatible with downstream enzymatic digestion and standard LC–MS/MS workflows. Conventional in-solution digestion with trypsin is limited by its incompatibility with strong detergents as they lack a dedicated detergent-removal step. Several strategies have been developed to address this limitation, including filter-aided sample preparation (FASP) ^19^, SP3 (Single-Pot, Solid-Phase-enhanced Sample Preparation) ^20^, and the Suspension-Trapping (S-Trap) platform ^21^, each offering different capabilities for processing detergent-containing samples. Advances in MS instrumentation and acquisition strategies have also improved proteomic profiling. Standard bottom-up shotgun proteomics involves enzymatic digestion followed by LC–MS/MS. Quantitative MS workflows are broadly categorized into isobaric labeling and label-free approaches. Isobaric tagging, exemplified by Tandem Mass Tags (TMT) ^22^, allows multiplexed analyses to reduce batch effects and enhance throughput. Label-free quantification using data-dependent acquisition (DDA) has traditionally been hindered by stochastic precursor selection, but this limitation has been addressed by data-independent acquisition (DIA) ^23, 24^, which systematically fragments all ions across the mass range to generate a comprehensive digital map of the proteome. When processed with dedicated algorithms (e.g., Spectronaut, DIA-NN) ^25^, DIA frequently exceeds the proteome coverage achieved by DDA, establishing it as a robust alternative to labeling-based workflows ^26^.

Using a comprehensive, multi-system evaluation of TurboID-based proximity proteomics in BV2 microglia in vitro and in astrocytes and neurons in vivo, we address key bottlenecks in protein solubilization, enrichment, digestion, and MS acquisition. We systematically compare three lysis chemistries, assess in-solution versus S-Trap digestion, evaluate on-versus off-bead proteolysis, benchmark DDA, DIA acquisition strategies as well as TMT labeling, assess the gains of newer and faster MS instruments (eg. ^27^, and examine the lower input limits of TurboID workflows using as little as 20 µg of lysate (as compared to >300 µg that has been used previously). We establish a workflow that is platform-independent and applicable to other biotin ligase systems (e.g., BioID2), extending its utility beyond the current study. Together, these optimizations yield a unified and scalable CIBOP workflow that increases proteome depth, improves recovery of synaptic and membrane-associated proteins, and generates high-quality datasets from limited material. This integrated pipeline enables reliable proteome mapping across diverse brain. ^28, 29, 30^

## RESULTS

### Optimization of TurboID Enrichment and Digestion Strategies in BV2 Microglia

Our technical objective was to optimize cell lysis conditions to maximize protein extraction, compare on-bead versus off-bead digestion protocols for streptavidin-enriched proteins, and evaluate the overall efficacy of S-Trap digestion ^21^ relative to traditional in-solution methods of digestion. To systematically benchmark these variables, we utilized a validated BV2 mouse microglial cell line stably expressing cytosolic V5–TurboID–NES ^9^. Following 24 hours of biotin supplementation, transduced and untransduced control cells (n = 2) were subjected to several parallel workflows (Fig. 1A). Four experiments compared streptavidin affinity purification (AP) under distinct lysis and digestion conditions: (1) 8 M urea lysis with in-solution digestion (ISD), (2) sequential 8 M urea/SDS lysis with S-Trap digestion, (3) 8 M urea lysis with S-Trap digestion, and (4) direct 5% SDS lysis with S-Trap digestion. For all S-Trap workflows, both on-bead and off-bead digestion strategies were evaluated. Additionally, we analyzed the total proteome (input) from 8 M urea and SDS lysates prior to enrichment to assess baseline coverage.

**Figure 1:**
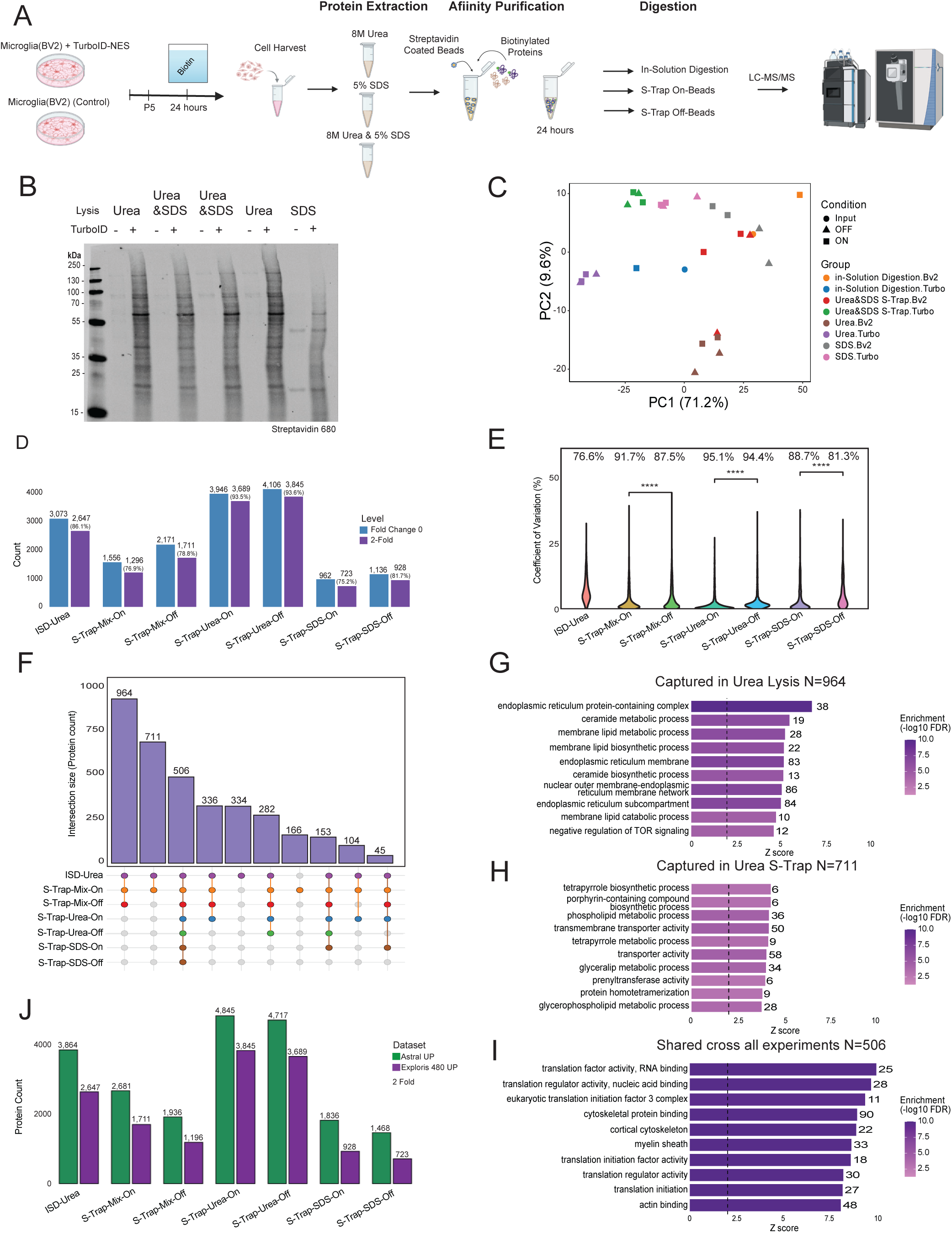
Optimization of BV2-TurboID proteomics workflow: **(A)** Experimental schematic illustrating BV2-TurboID sample preparation across multiple lysis chemistries followed by streptavidin enrichment, on-bead in-solution digestion, and LC-MS/MS analysis. **(B) Western blot with** SDS showing increased biotin labeled protein abundance in TurboID across lysis conditions, confirming successful proximity labeling. **(C)** PCA of BV2 proteomes generated from all conditions (PC1 = 71.2%, PC2 = 9.6%; axis variance values indicated) highlights workflow-dependent clustering, with Urea, Urea+SDS and SDS S-Trap preparations separating distinctly from in-solution digestion. **(D)** Protein identification counts at baseline (FC=0) and under ≥2-fold filters demonstrate highest proteome depth in S-Trap-Urea and S-Trap-Mix TurboID-ON preparations (up to 4,106 proteins), markedly exceeding ISD-Urea and SDS protocols. **(E)** Coefficient of variation distributions shows lower variability in S-Trap workflows compared to ISD-Urea, indicating improved quantification reproducibility. Violin plot showing the distribution of protein-level coefficient of variation (CV%) across BV2 TurboID sample preparation workflows. S-Trap–based protocols exhibited markedly reduced variability relative to ISD-Urea digestion, with S-Trap-Mix-ON, S-Trap-Urea-ON, and S-Trap-SDS-ON producing tighter CV distributions (****p < 0.0001 for each indicated comparison), indicating superior quantitative reproducibility under TurboID-ON conditions. The x-axis denotes lysis and capture strategy, and the y-axis displays total CV% relative to replicate intensity variance. **(F)** Intersection analysis (FC>= 2) reveals large, shared protein cores across S-Trap strategies (964– 711 proteins per intersection) with smaller method-specific sets, suggesting strong retention of biologically relevant targets under optimized workflows. **(G,H,I)** GO term enrichment identifies lysis-specific biological biases: Urea preferentially captures ER-lipid metabolic machinery; Urea+SDS S-Trap enriches tetrapyrrole, transporter and lipid pathways; and shared proteins across all workflows map to translation regulators, cytoskeleton and myelin-associated complexes (x-axis = Z-score as enrichment ; color scale = -log10 FDR). **(J)** Comparison of upregulated proteins between BV2 Astral and BV2 Exploris480 instruments shows consistently greater protein recovery on Exploris480 across all lysis-S-Trap configurations (4,845 vs 3,864; 4,717 vs 3,845; etc.), validating robustness and sensitivity of the selected LC-MS platform.

We first validated streptavidin AP efficiency by gel electrophoresis, confirming a marked increase in total protein captured from TurboID-expressing cells. Western blot analysis further verified the specific enrichment of biotinylated proteins (Fig. 1B). Following AP, we quantified peptide recovery after digestion. Despite using approximately 90% of the captured beads for the ISD workflow (compared to 45% for each S-Trap experiment), the peptide yield from ISD was 10-fold lower than from the S-Trap workflows (Supp Table. 1). This discrepancy highlights a major inefficiency in peptide elution and/or digestion within the ISD workflow, suggesting that MS workflows that incorporate S-Trap rather than ISD should yield deeper proteome coverage.

### S-Trap Digestion Improves Quantitative Depth and Reproducibility of Microglia Proteome

DIA-MS analysis ^24, 25^ of all samples identified 6,335 total proteins within input and affinity-purified (AP) fractions. Comparison of enrichment workflows revealed that 8 M urea lysis coupled with S-Trap digestion provided the greatest depth; the off-bead and on-bead protocols identified 3,897 and 3,721 proteins in TurboID-positive samples, respectively, compared to 1,237 and 915 in controls. While the 8 M urea/ISD workflow identified fewer proteins (2,605), it exhibited the lowest background (57 proteins in controls). SDS-based protocols yielded mixed results: direct 5% SDS lysis captured ∼1,000 proteins (on-and off-bead), whereas sequential urea/SDS lysis recovered 2,110 (off-bead) and 1,579 (on-bead) proteins. To verify that these differences stemmed from enrichment efficiency rather than bias due to lysis conditions, we profiled the input (whole cell) fractions. We confirmed deep and comparable proteome coverage across S-Trap 5,897 proteins and ISD with 5,588 proteins (with 84.5% [n=5,260] overlap). This indicates that the yield of the S-trap digestion reflects enhanced recovery of biotinylated proteins rather than differences in starting material (Supp Table. 1).

The in-solution digestion workflow exhibited a lower background, with the fewest proteins identified in control samples. We attribute this to a technical artifact of the injection protocol rather than true specificity. As shown by peptide-BCA quantification, ISD recovered ∼10-fold less total peptide mass despite starting with more captured protein (Supp Table. 1). Thus, fixed-volume injections for ISD control samples contained substantially less material, artificially suppressing the background (control) and inflating the apparent signal-to-noise ratio. This likely reflects under-sampling of contaminant proteins rather than a genuine reduction in non-specific binding.

To visualize global variance across workflows, we performed principal component analysis (PCA) on enriched proteins (FC ≥ 2 in AP samples from TurboID compared to Control BV2 cells). Sample groups were clearly separated primarily by lysis buffer, with urea-based and SDS-containing lysates forming distinct clusters, suggesting lysis buffer is the dominant determinant of proteomic variance in this system (Fig. 1C). Within these clusters, replicates for urea-based workflows were grouped closer together, consistent with reproducible extraction of the microglial proteome under these conditions.

Next, we assessed the reproducibility of our results by calculating the coefficient of variation (CV) for each workflow using 2 technical replicates per sample. All S-Trap–based approaches demonstrate excellent quantitative precision with on-bead digestion consistently yielding lower variance than off-bead digestion. In urea lysis S-Trap on-bead CV was 2.11% versus 2.80% off-bead, and in SDS lysis S-Trap on-bead CV was 3.52% versus 5.51% off-bead. The on-bead 8 M urea S-Trap workflow exhibited the highest overall reproducibility (CV = 2.11%) (Fig. 1E). We next analyzed enriched proteins within each workflow to identify proximity-labeled proteins based on enrichment in TurboID samples relative to controls.

We next identified proximity-labeled proteins based on ≥2-fold enrichment in TurboID samples relative to controls. Urea lysis with off-bead S-Trap digestion yielded the greatest depth of 4,106 proteins, with 3,845 (93.6%) retained after filtering. ISD with urea lysis identified 3,072 proteins, of which 2,647 (86.1%) remained. Mixed lysis retained 1,711 of 2,171 proteins (78.8%) with off-bead digestion and 1,296 of 1,556 proteins (76.9%) with on-bead digestion. SDS lysis produced fewer enriched proteins, retaining 928 of 1,136 (81.7%) with off-bead digestion and 723 of 962 (75.2%) with on-bead digestion (Fig. 1D). Across all lysis conditions, off-bead digestion consistently resulted in higher protein retention after application of the fold-change cutoff.

### On-Bead Digestion Captures More Unique Proteins than Off-Bead Digestion in Microglial BV2 cells

We next examined the technical variable of on-bead versus off-bead digestion. On-bead digestion is sometimes avoided due to co-elution of streptavidin peptides that dominate DDA-MS spectra, but the comprehensive fragmentation of DIA-MS may mitigate this issue. A direct comparison showed that on-bead digestion consistently identified more unique proteins, with 2,150 unique proteins compared with only 172 unique proteins in the off-bead workflow, with 1,538 proteins shared (Supplementary Fig. 1B). Of the 3,860 proteins identified across the urea S-Trap conditions, 56% were unique to the on-bead workflow, while only 4% were unique to off-bead. Functional annotations indicated that shared proteins were enriched for fundamental processes such as translation and vesicle transport, while on-bead–unique proteins were enriched in dynamic and regulatory protein pathways like RNA processing, autophagy, and intracellular trafficking that may have lost in elution-dependent off-bead protocols (Supplementary Fig. 1C, D, and E). Taken together, these results indicate that 8 M urea lysis with S-Trap digestion achieved the greatest proteome depth in vitro. Within this workflow, on-bead digestion yielded a larger fraction of uniquely identified proteins, whereas off-bead digestion showed improved reproducibility and higher protein retention after fold-change filtering

An UpSet plot of highly-enriched proteins (FC ≥ 2) revealed a large core group of proteins shared across on- and off-bead workflows (Fig. 1F), although the superior depth of the on-bead method remained evident. To compare biological signatures, we examined the three largest intersections: the universal core proteome 506 proteins, the 711 proteins enriched exclusively by 8 M urea lysis with S-Trap workflows, and the 964 proteins shared between 8 M urea lysis followed by ISD, and S-Trap workflows (Fig. 1F). GO analysis of these groups further revealed distinct enrichment patterns with core group enriched basic housekeeping functions such as cytoskeletal binding and translation initiation (Fig. 1I). The 8 M Urea lysis with S-Trap– exclusive group was enriched for phospholipid and glycolipid metabolism (Fig. 1H), and the 8 M urea lysis core proteome was enriched for ER membrane functions and ceramide metabolism (Fig. 1G). These findings demonstrate that while a core proteome is consistently detected, the optimized S-Trap workflow provides deeper access to specific metabolic pathways due to changes in the accessibility of membrane associated proteins.

### DIA and the Astral Platform Substantially Expand BV2 Proteome Coverage

DIA-MS data from two Orbitrap MS from Thermo Fisher Scientific were compared. The Astral, which employs a new analyzer for MS2 in parallel with the Orbitrap analyzer for MS1, provides substantially increased scan speed and ion transmission efficiency compared with the Exploris 480^27^, making it particularly well-suited for the low-abundance, complex peptide mixtures typical of proximity-labeling experiments. The Astral MS consistently outperformed the Exploris 480 MS across all BV2 workflows. For every experimental condition, the Astral identified more labeled proteins than the Exploris 480 at the same enrichment thresholds. The largest gains were observed in the high-performing 8 M urea lysis, S-Trap workflows, where Astral detected 4,845 and 4,717 proteins with on-and off-bead digestion respectively, compared with 3,845 and 3,689 proteins on the Exploris 480, representing a nearly 30% increase in proteome depth. Even in lower-yield experimental conditions, Astral retained a clear advantage. These data demonstrate that the increased sequencing speed and sensitivity of the Astral substantially deepens TurboID proteome coverage. The main conclusions regarding which sample-preparation pipeline (lysis, digestion, enrichment) are optimal remained unchanged regardless of whether data were acquired on Exploris 480 or Astral (Fig. 1J).

### Optimization of the CIBOP Workflow Expands the Microglial Proteome Compared to Previous Datasets

To assess the proportion of the whole-cell proteome captured by TurboID, we compared the 8 M urea lysis/S-Trap on-bead dataset against the total input proteome. Analysis of replicates identified 3,896 proteins, whereas the input proteome yielded 5,587 proteins via ISD and 5,896 via S-Trap. This corresponds to a coverage of 69.7% relative to the ISD input and 66.1% relative to the S-Trap input. These values represent an improvement over our previous BV2 study^14^, where the TurboID-enriched proteome encompassed approximately 59% of the detectable whole-cell proteome. This increased depth suggests that the optimized S-Trap workflow captures a more physiologically representative fraction of the total cellular proteome.

To benchmark the proteomic depth of our optimized workflow against our previously published BV2–TurboID study^14^, we compared unique protein identifications from the two datasets. The earlier study, which employed 8 M urea lysis followed by in-solution digestion (ISD) with DDA-MS, identified 2,163 unique proteins (2,183 total). In contrast, our current 8 M urea on-bead S-Trap DIA-MS workflow identified 4,015 unique proteins (4,016 total). Combining the two datasets generated a library of 4,691 distinct proteins. The optimized method identified 2,528 proteins that were not detected in the earlier study, whereas only 676 proteins were unique to Sunna et al. (Supplementary Fig. 1F). Thus, 54% of the combined protein pool was exclusive to the new S-Trap workflow, compared to only 14% exclusive to the ISD workflow, with 32% being shared.

GO enrichment analysis revealed that proteins uniquely recovered by the optimized on-bead S-Trap workflow were strongly enriched for cytoplasmic translation, ribonucleoprotein complexes, cytosolic ribosomes / ribosomal subunits, and polysomes, indicating improved recovery of core translational machinery (Supplementary Fig. 1G). When we examined a broader, mixed set of proteins captured across both workflows, we observed prominent enrichment of regulatory and signaling categories, including kinase activity, protein phosphorylation, transmembrane signaling receptor activity, lipid transporter activity, and phospholipid and glycerolipid metabolic processes, highlighting enhanced sensitivity for low-abundance signaling and lipid-regulatory proteins (Supplementary Fig. 1H). In contrast, proteins uniquely identified in our prior ISD workflow^14^ were predominantly enriched for aerobic respiration, cellular respiration, tricarboxylic acid cycle, oxidative phosphorylation, and the aerobic electron transport chain, consistent with a stronger bias toward mitochondrial energy metabolism (Supplementary Fig. 1I). Together, these data indicate that while the reference method favors recovery of core mitochondrial metabolic machinery, the optimized workflow preferentially enhances detection of protein translation components and regulatory signaling pathways, offering a more functionally diversified view of the proteome.

### Optimization of Astrocyte CIBOP Workflow in vivo

Astrocytes are critical cellular components of the central nervous system, with key roles in maintaining ion homeostasis, recycling neurotransmitters, and supporting synaptic function ^1, 31, 32, 33^. They are also implicated in numerous neurodegenerative diseases, including Alzheimer’s disease, Parkinson’s disease, and Huntington’s disease ^2, 28, 34, 35, 36, 37, 38, 39^. To evaluate TurboID workflow performance in vivo, we performed technical optimization experiments using brain tissue from mice in which the CIBOP strategy was used to label astrocyte proteomes (Aldh1l1-CreERT2/TurboID)^11^.

We generated these mice by crossing a floxed, V5-tagged TurboID mouse line (Rosa26-TurboID-V5-NES) with the tamoxifen-inducible Aldh1l1-CreERT2 driver line. In the resulting offspring (heterozygous for Cre and TurboID), administration of tamoxifen induces Cre recombination leading to expression of a cytosolic V5–TurboID enzyme and proteomic labeling specifically in Aldh1l1-positive astrocytes in the nervous system^11^. To directly compare lysis methods and determine whether in vivo outcomes differ from our in vitro benchmarking, individual mouse brains were bisected into hemispheres. One hemisphere underwent urea-based lysis and the other SDS-based lysis, enabling a paired comparison of workflow performance using tissue from the same animal (Fig. 2A).

**Figure 2:**
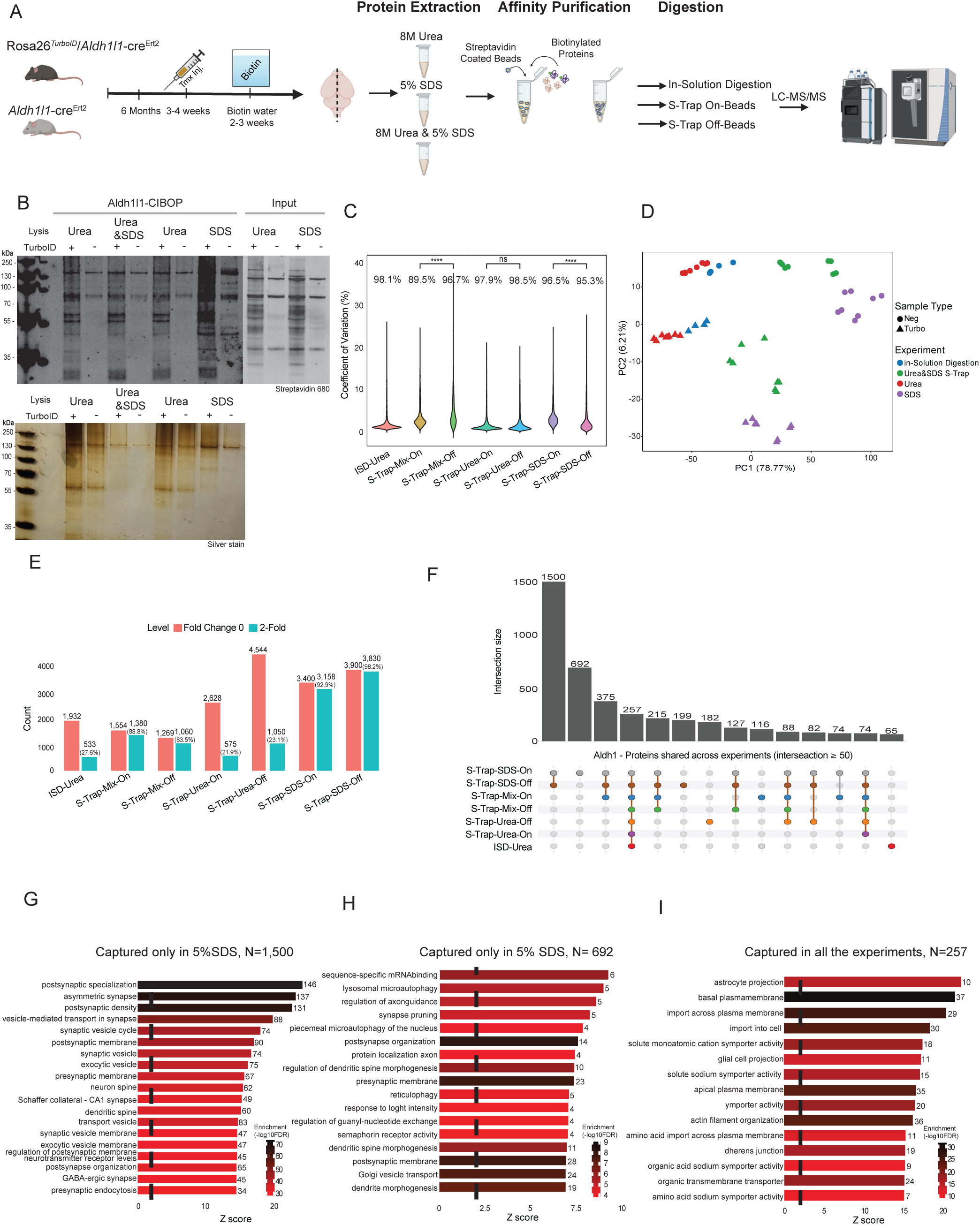
Optimization of Aldh1l1-TurboID astrocyte proteomics workflow. **(A) Experimental workflow schematic:** Mouse models showing Rosa26^TurboID/+ Aldh1l1-cre^ERT2 and control, tamoxifen induction, biotin supplementation, brain dissection, protein extraction using Urea/SDS conditions, streptavidin pulldown, digestion via in-solution or S-trap (on/off bead), followed by LC-MS/MS analysis. **(B) Western blot and silver stain evaluation:** Comparison of biotinylated protein enrichment under Urea, Urea+SDS, and SDS extraction conditions. TurboID+ samples show strong band recovery and clear enrichment relative to TurboID controls, confirming successful labeling and pulldown efficiency. **(C)CV (Coefficient of Variation):** Violin plots showing reproducibility across digestion strategies. S-Trap Urea and S-Trap SDS conditions exhibit lower variability than in-solution digestion. Statistical comparisons indicate significant improvement in consistency for on-bead S-Trap workflows. **(D) Aldh1 DDA PCA:** Principal Component Analysis (PC1 = 78.77%, PC2 = 9.21%) reveals clear separation between TurboID positive and TurboID negative samples, with grouping based on extraction methods (Urea, SDS, Urea+SDS S-Trap, in-solution). Sample type clustering confirms workflow-driven proteomic differences. **(E) Protein counts comparison across experiments:** Bar plot showing total identified proteins (Fold Change 0) and proteins increased ≥2-fold across experiments. S-Trap SDS Off and S-Trap–SDS-On yield the highest recovery, followed by S-Trap-Urea-On. Percentage values represent a proportion of 2-fold proteins relative to the total identified. **(F) Aldh1 Proteins shared across experiments (intersection** ≥ **50):** UpSet plot showing intersection size of proteins detected across extraction workflows (S-Trap-SDS-On, S-Trap-SDS-Off, S-Trap-Mix-On, S-Trap-Mix-Off, S-Trap-Urea-On, S-Trap-Urea-Off, ISD-Urea). The largest shared set contains 1500 proteins, followed by 692, 375, 257, 215, 199, and smaller intersections. Each connected dot set represents a specific workflow combination contributing to that shared protein group.**(G) GO enrichment for captured only in 5% SDS with S (Upset 1500):** Enriched terms include postsynaptic specialization, asymmetric synapse, synaptic vesicle transport/cycle, neuron spine, dendritic spine, and synaptic membrane organization. **(H) GO enrichment for captured only in 5% SDS S (Upset 692):** Enriched for mRNA binding, lysosomal micro-autophagy, axon guidance, dendritic spine morphogenesis, synapse pruning. **(I) GO enrichment for captured in all experiments (Upset 257):** Shared core pathways include astrocyte projection, membrane transport/import, ion symporter activity, glial projections, and amino acid transport. Z-scores reflect enrichment; color scale indicates −log10(FDR).

We validated streptavidin AP efficiency by silver staining of eluates resolved by SDS–PAGE, confirming substantially greater biotin-enriched protein capture in Aldh1l1–TurboID samples compared with controls. Western blot and silver stain analysis further verified the specific enrichment of biotinylated proteins (Fig. 2B). Consistent with our in vitro results, peptide recovery following S-Trap digestion was 10- to 20-fold higher relative to ISD even when 45% of the beads used for S-Trap digestion and 90% for ISD after streptavidin AP, highlighting the same digestion efficiency advantage seen in BV2 cells (Supp Table. 2).

DIA-MS analysis identified 7,766 proteins across all input and affinity-purified (AP) conditions. Among the enrichment strategies, 8 M urea lysis yielded the highest number of identifications in Aldh1l1-TurboID-positive samples, with the ISD workflow identifying 6,316 proteins, followed by S-Trap off-bead (6,277) and on-bead (5,936) protocols. However, urea-based conditions exhibited substantial nonspecific binding, as untransduced controls yielded similarly high identification rates (5,993 for ISD, 5,894 for off-bead, and 5,499 for on-bead). In contrast, while SDS lysis resulted in lower total identifications in TurboID samples (4,726 off-bead; 4,166 on-bead), it demonstrated a significant reduction in background; untransduced controls in SDS conditions yielded 2,757 and 2,215 proteins for off-bead and on-bead workflows, respectively.

We quantified reproducibility by calculating the coefficient of variation (CV) across the experimental compared samples. The urea-based S-Trap workflows exhibited the lowest overall variance, with CVs of 1.6% for on-bead digestion and 1.7% for off-bead digestion, indicating that urea lysates remain technically stable regardless of digestion format. In contrast, the digestion strategy strongly influenced variability within SDS-based workflows. The SDS on-bead protocol yielded a median CV of 2.6%, significantly lower than the 3.6% observed for off-bead digestion (p < 0.0001). This divergence suggests that the additional elution and transfer steps required for off-bead digestion introduce stochastic variation when detergents are present, whereas on-bead digestion minimizes this technical noise (Fig. 2C).

We used PCA on proteomic data to visualize global variance across the in vivo workflows. The first principal component (PC1), explaining 78.8% of the variance, separated samples based on urea lysis and SDS-based lysis methods while PC2 (6.2% of the variance), distinguished biotin-enriched TurboID samples from non-biotinylated controls. Notably, the SDS-based S-Trap workflow exhibited the most robust separation of TurboID versus control samples along PC2, consistent with its high enrichment efficiency (Fig. 2D). In contrast, urea-based workflow showed less separation compared to SDS. A more detailed PCA separating on-bead and off-bead conditions is provided in (Supplementary Fig. 2A).

To determine the optimal workflow for Aldh1l1–TurboID brain tissue, we first compared the total number of statistically significant proteins identified by each protocol using a permissive threshold (p ≤ 0.05 comparing TurboID to non-TurboID control groups). The urea-based S-Trap workflows yielded the highest number of enriched proteins: the 8 M urea lysis, off-bead digestion identified 4,544 proteins, and the on-bead group identified 2,628 proteins. The ISD workflow identified 1,932 proteins. SDS-containing workflows also performed well, with direct SDS lysis identifying 3,900 proteins for off-bead and 3,400 proteins for on-bead, while sequential 8 urea/SDS lysis yielded 1,269 proteins for off-beads and 1,554 proteins for on-bead digestion groups. The influence of lysis buffer became evident only after applying a more stringent enrichment threshold (FC ≥ 2, p ≤ 0.05), which isolates high-confidence, strongly enriched DEPs. Under this cutoff, SDS-containing workflows far outperformed urea-based protocols. The SDS lysis off-bead SDS workflow identified 3,830 DEPs (retaining 98.2%) of its initial hits, while the SDS lysis on-bead workflow identified 3,158 DEPs (retaining 92.9%). Sequential urea/SDS lysis also performed well, yielding 1,380 DEPs for on-bead digestion (retaining 88.8%) and 1,060 DEPs for off-bead digestion (retaining 83.5%). In contrast, workflows relying exclusively on urea exhibited steep attrition. The urea lysis off-bead workflow identified only 1,050 DEPs (retaining 23.1%), and the corresponding on-bead variant yielded only 575 DEPs (retaining 21.9%). The ISD workflow identified just 533 DEPs, retaining only 27.6% of its initial hits. These patterns demonstrate that although 8 M urea-based lysis may produce large number of enriched proteins compared to non-labeled controls, the majority of these do not meet 2-fold enrichment thresholds. SDS-based workflows, by contrast, provide deeper access to the strongly enriched fraction of the astrocyte proteome and exhibit superior signal-to-noise after stringent cutoff application (Fig. 2E).

### SDS Lysis Extends Astrocyte CIBOP Proteomes via Labeling of Membrane-Rich and Synapse-associated Proteins

Given the greater astrocytic proteomic coverage using SDS-based workflows, we further analyzed the overlap between the on-bead and off-bead digestion protocols (Supplementary Fig. 2B). We observed a high degree of concordance, with 2,896 proteins shared between the two strategies. The off-bead protocol uniquely identified 933 proteins, whereas the on-bead workflow uniquely captured 262 proteins. GO enrichment of the shared proteome (2,896) (using 4,091 proteins as background list) revealed strong overrepresentation of tripartite synapse–associated terms, including postsynaptic specialization, asymmetric synapse, postsynaptic density, and synaptic vesicle cycle (Supplementary Fig. 2C). These results indicate that SDS lysis, as compared to urea lysis, may more effectively solubilize dense synaptic structures regardless of the subsequent digestion strategy.

Proteins unique to the off-bead fraction (933) were predominantly associated with intracellular trafficking and cellular organization, including endosomal transport, Golgi vesicle transport, and post-synapse organization (Supplementary Fig. 2D). In contrast, the smaller set unique to the on-bead fraction (262) displayed enrichment for regulatory processes such as negative regulation of RNA splicing and calcium-dependent protein binding (Supplementary Fig. 2E). Together, these findings suggest that while the off-bead workflow captures a slightly broader set of trafficking-related proteins, the on-bead SDS workflow captures the vast majority of the core synaptic structure 2,896 proteins with higher reproducibility.

To further evaluate the distribution of differentially enriched proteins (DEPs) across workflows, we assessed overlapping and unique features of the proteomes across all experimental workflows (Upset R plot, Fig. 2F). This analysis revealed that the SDS-lysis workflows were by far the most effective, capturing the largest number of proteins across all conditions. The largest intersection contained 1,500 proteins identified in both S-Trap on- and off-bead digestion workflows and not detected in other conditions. The second-largest intersection of 692 proteins was unique to the SDS lysis off-bead workflow alone. The core proteome shared across all workflows was 257 proteins, while proteins unique to urea-based workflow in vivo, were far less than those observed in vitro (Fig 1.F). This discrepancy may be explained by differences in tissue complexities (brain versus cell lines in culture) and the ability of SDS to better solubilize membrane-associated proteins.

To understand the biological relevance of the proteins captured by the different in vivo workflows, we performed a GO analysis on the major intersections identified (Fig. 2F). The core proteome of 257 proteins was enriched for astrocyte projection, glial cell projection, and membrane transporter/ion homeostasis pathways, consistent with canonical astrocytic structural and transport functions. In contrast, the 1,500 proteins quantified by on-bead and off-bead SDS-based S-Trap workflows demonstrated strong enrichment for synaptic GO terms, including postsynaptic density, the synaptic vesicle cycle, and the Schaffer collateral CA1 synapse. This pattern suggests that SDS lysis substantially improves recovery of membrane-proximal and synaptic components of astrocytes, particularly those situated at tripartite synapses. The SDS off-bead workflow additionally recovered 692 proteins whose GO signatures prominently featured post-synapse organization, presynaptic membrane organization, and regulation of dendritic spine morphogenesis, further supporting enhanced capture of peri-synaptic astrocyte processes and their immediately apposed neuronal compartments.

To determine whether synaptic GO signatures in our SDS-based proteomes reflect true perisynaptic astrocyte biology rather than neuronal contamination, we examined representative proteins underlying postsynaptic density associated GO terms using published cell-type–resolved transcriptomic and proteomic datasets. Several proteins consistently detected in these synaptic categories including Stat3, Ntrk2 (TrkB-T1), Adam22, Ptk2b, Tsc1/Tsc2, Rheb, and Prnp ^40, 41, 42, 43, 44^ which have demonstrated expression in astrocytes and are consistent with astrocyte signaling, morphology, or membrane-associated functions. Conversely, classical neuronal scaffold and receptor proteins such as Dlg4 (PSD-95), Shank3, Homer2, Gria1/2/3, and Rims1 ^45, 46, 475, 46, 47^ were also captured in astrocyte proteomes, consistent with labeling of neuronal synaptic membranes which are juxtaposed to astrocyte processes. A subset of proteins, including Slc8a1, Snx27, Vps35, Rab8a, and Clstn1, are expressed in both neurons and astrocytes and likely represent shared trafficking and membrane-interaction machinery at the synaptic interface ^48, 49, 50^. Together, these mixed expression profiles indicate that astrocyte–TurboID labeling captures a combination of astrocyte-enriched, neuron-enriched, and shared/interface proteins. This pattern is consistent with the nanoscale spatial organization of tripartite synapses, wherein astrocytic processes physically envelop and regulate neuronal synaptic compartments ^32, 33^ (Fig. 2G, H, I)

Unlike results from BV2 microglia in vitro, in vivo astrocyte CIBOP proteomes were best captured when SDS lysis was incorporated, suggesting that tissue-specific matrix effects and the dense, membrane-structure of astrocytes in vivo drive differential extraction efficiency between buffer conditions. We therefore compared the SDS and urea input lysates, and revealed an extraction advantage for SDS brain lysates. Across 7,577 proteins identified at the input level, 4,070 exhibited higher abundance in SDS lysates, whereas 3,430 were more abundant in urea with normalization. Together, these data confirm that SDS provides better solubilization of membrane-rich and synaptic compartments relative to urea and enhances recovery of biologically relevant proteins at the input level. While the shared protein fraction contained numerous canonical astrocytic markers and core functional proteins, we next examined how newly identified proteins broadened our understanding of astrocyte biology. GO analysis of the 2,280 proteins uniquely identified by our optimized SDS-S-Trap DIA workflow revealed strong enrichment for RNA processing, mRNA metabolic processes, and transcriptional regulation (Fig. 3B). These findings indicate that our optimized workflow improves solubilization of complex subcellular compartments and enhances sensitivity toward lower-abundance regulatory components that modulate astrocyte gene expression and cellular state. The 878 proteins detected by both our workflow and the previously published Urea-ISD-DDA method were enriched for basal and basolateral plasma membrane, cytoskeletal and somatodendritic structures (Fig. 3C), representing abundant, well-established features of astrocyte biology that serve as functional anchors within the shared dataset. The 427 proteins uniquely captured by the prior Urea-ISD-DDA workflow from Rayaprolu et al.^11^ were predominantly enriched for mitochondrial metabolic pathways, including carboxylic acid, amino acid, and fatty acid metabolic processes, oxidoreductase activity, and aerobic electron transport chain components (Fig. 3D). This pattern reflects a workflow bias toward highly abundant metabolic machinery. Together, these results demonstrate that our optimized SDS-S-Trap DIA workflow extends the accessible astrocytic proteome beyond structural and metabolic constituents, revealing regulatory pathways and nuclear processes that are largely missed by standard approaches, and thereby providing a more functionally informative and comprehensive view of astrocyte biology.

**Figure 3:**
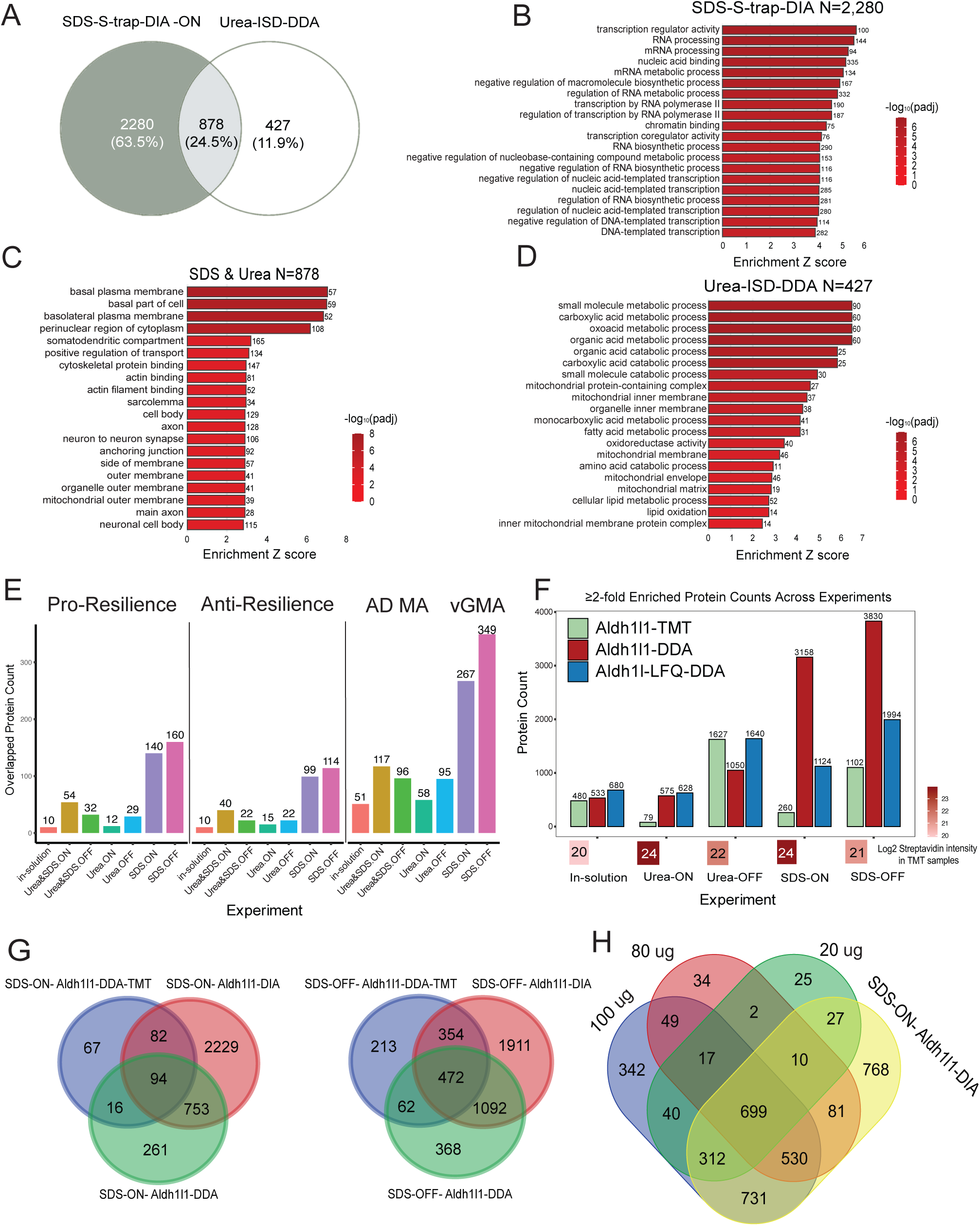
Comparative evaluation of Aldh1l1-TurboID astrocyte proteomics across datasets, quantification strategies, and input amounts: **(A) Comparative analysis of Rayaprolu dataset vs. our DIA dataset:** Venn diagram illustrates the overlap between proteins identified in the Rayaprolu SDS-S-trap DIA ON-bead dataset and our Urea-ISD DIA (OFF-bead) dataset. A total of 2,280 proteins (63.5%) were uniquely detected in the Rayaprolu dataset; 427 proteins (11.9%) were unique to our DIA dataset, and 878 proteins (24.5%) were common to both datasets. **(B) GO enrichment for upregulated 2280 proteins (Rayaprolu et al unique proteins):** These proteins show strong enrichment for RNA metabolism and processing pathways, including transcription regulator activity, RNA and mRNA processing, nucleic acid binding, regulation of macromolecule biosynthesis, RNA polymerase II transcription, and chromatin binding. These terms highlight a broad enrichment of nuclear and transcriptional regulatory functions. **(C) GO enrichment for 878 Shared proteins between Rayaprolu and DIA:** The 878 overlapping proteins are significantly enriched for neuronal, synaptic, and cytoskeletal compartments, including basal plasma membrane, perinuclear region, somato dendritic compartment, cytoskeletal protein binding, actin and axon-related structures, anchoring junctions, and neuronal cell body components. These shared proteins capture core neuronal architecture and transport functions common across both datasets. **(D) GO enrichment for upregulated 427 proteins:** The 427 proteins detected uniquely in DIA are enriched for mitochondrial and metabolic pathways, including top enrichment for small molecule metabolic processes, carboxylic acid and organic acid metabolism, mitochondrial inner membrane, protein- containing complex. These enrichments indicate that DIA method captured deeper mitochondrial and metabolic protein coverage relative to the Rayaprolu et al dataset. **(E): Differentially enriched proteins (DEPs) for Aldh1l1 across methods reveal enrichment of neurological disorder linked resilience genes:** Stacked bar charts display the fraction of overlapping proteins across experiments in Aldh1l1 proteomics, categorized into Pro-resilience, Anti-resilience, and AD MAGMA (genetic risk^51^) proteins sets. Across all in-solution, Urea ON/OFF, and SDS ON/OFF shows a consistent enrichment of AD MAGMA associated proteins (blue) is observed, ranging from 55–80% of overlapping proteins. Pro-resilience proteins (green) and Anti-resilience proteins (orange) are also represented, with method-specific differences. SDS-OFF and SDS-ON chemistries show the highest counts of resilience-related proteins, reflecting deeper proteomic capture of disease-relevant pathways. **(F): Comparison of TMT, LFQ, and DIA workflows shows highest protein recovery with SDS-based DIA:** Barplot compare protein counts across in-solution, Urea ON/OFF, and SDS ON/OFF digestions for Aldh1-TMT, Aldh1-DDA, and LFQ-DDA datasets. SDS-based preparations consistently yield the highest number of ≥2-fold proteins, with SDS-OFF showing the greatest recovery for Aldh1-DDA (3,830 proteins) and strong performance in LFQ-DDA (1,994 proteins). Urea-OFF conditions also show robust recovery across all methods, particularly in LFQ-DDA (1,640 proteins). In contrast, Urea-ON yields the lowest TMT recovery (79 proteins). The accompanying heatmap summarizes the expression of Streptavidin in Aldh1-TMT across all positive Turbo samples within each experiment. **(G) Venn comparison of TMT, LFQ-DDA, and DIA shows DIA captures the largest unique proteome with a consistent core shared across methods:** SDS-ON and SDS- OFF Venn diagrams highlight DIA’s high sensitivity (SDS-ON: 2,229 DIA-only; SDS-OFF: 1,911 DIA-only) and a conserved overlap across experiments (SDS-ON: 94 shared; SDS-OFF: 472 shared). LFQ-DDA and DIA show strong overlap, while TMT contributes to smaller but reproducible unique sets. SDS-OFF yields the deepest intersection across all quantification methods. **(H) Venn comparison of low-input DIA (20, 80 and 100 µg) shows a large, shared proteome with strong recovery even at 20 µg:** Across 100 µg, 80 µg, 20 µg, and SDS-ON DIA, a robust core of 699 proteins is consistently detected. Higher inputs yield more unique proteins, but 20 µg still retains substantial overlap, demonstrating strong DIA sensitivity at low sample amounts.

To benchmark the depth and biological specificity of our in vivo astrocyte proteome, we compared our dataset to the landmark study by Rayaprolu et al.^11^, which identified 1,305 enriched astrocyte proteins using an affinity purification–based strategy. In contrast, our optimized SDS lysis combined with on-bead S-Trap digestion and DIA-MS identified 3,158 differentially enriched proteins from Aldh1l1–TurboID mice. When the two datasets were compared directly, 878 proteins overlapped, whereas 2,280 proteins were uniquely recovered by our workflow and 427 were unique to the published dataset (Fig. 3A). This substantial increase in proteome depth demonstrates that SDS-based lysis, coupled with the enhanced sensitivity of DIA, provides access to a significantly expanded astrocyte proteome.

To evaluate how methodological improvements influenced recovery of Alzheimer’s disease– relevant proteins, we interrogated astrocyte-TurboID proteomes generated using different workflows for known AD genetic risk factors (AD MAGMA gene lists ^51^) and proteins associated with cognitive resilience in humans ^52^ (Fig. 3E). Astrocyte proteomes obtained using SDS lysis coupled with the S-Trap workflow yielded the highest number of both AD MAGMA and resilience-associated proteins across all experiments. This reflects the substantially greater proteome depth achieved with the optimized workflow, which in turn enables more comprehensive capture of astrocyte pathways implicated in AD susceptibility as well as resilience. Together, these data demonstrate that improving proteome recovery directly enhances detection of disease-relevant and protective protein networks in astrocytes. The off-bead SDS protocol provides access to the largest number of unique astrocyte-enriched proteins.

### Suitability of the Optimized Workflow for Tandem Mass Tag and Low Input Proteomics

To rigorously validate our acquisition strategy, we benchmarked the optimized SDS–S-Trap– DIA workflow against conventional data-dependent acquisition (DDA) and multiplexed 10-plex TMT isobaric labeling using identical starting biological material (n = 10: ISD, Urea, SDS lysated with On&Off-Beads with S-Trap digestion) (Fig.3F) ^22^. We first compared performance across the label-free modes. In the DDA dataset, the urea-based off-bead protocol yielded the highest number of enriched proteins (1,627). This is consistent with the expectation that peptide elution (Off-beads) prior to digestion enhances peptide accessibility for precursor selection in DDA acquisition mode. Applying the optimized SDS–DIA workflow resulted in a substantially deeper proteome, identifying 3,158 enriched proteins. Importantly, this depth was achieved under a stringent statistical filter (p < 0.05 and FC > 2 across n = 4 biological replicates). These results demonstrate that the increased sensitivity and systematic sampling afforded by DIA acquisition surpass the accessibility benefits of off-bead DDA workflows^12, 24^, and that the use of SDS-based lysis further enhances recovery of hydrophobic and synaptic proteins compared with urea-based methods ^16^.

We next evaluated the compatibility of the optimized workflow with multiplexing using a TMT- 10plex strategy. Consistent with label-free trends, on-bead digestion produced fewer identifications than off-bead digestion, but this reduction was exacerbated in the TMT setting. While contaminant streptavidin from the beads contributes to this loss, our analysis suggests a specific interference mechanism: in S-Trap on-bead samples, we observed that abundant streptavidin peptides were extensively labeled by the TMT reagents. This high load of non-target contaminants affects the TMT label, competitively reducing the labeling efficiency for the target peptides. Furthermore, these labeled streptavidin peptides dominate the reporter ion channels, leading to severe reporter ion compression and compromised quantification (Fig 3F). Consequently, off-bead digestion is strongly recommended when applying TMT multiplexing to remove streptavidin prior to labeling.

By contrast, DIA-based LFQ quantification preserved accurate measurement across a wide dynamic range, enabling confident quantification of 3,158 enriched proteins (Fig. 3F). Across

both ON- and OFF-bead conditions, DIA consistently recovered the largest set of uniquely identified streptavidin-enriched proteins 2,229 and 1,911, respectively far exceeding those detected by DDA-LFQ or DDA-TMT (Fig. 3G). Only a small core set of proteins was shared across all three acquisition strategies (94 ON-bead and 472 OFF-bead), indicating that DIA substantially expands accessible proteome coverage beyond the limited subset captured by traditional DDA approaches. Collectively, these comparisons indicate that while DDA and TMT methods have specific strengths, the optimized SDS–DIA workflow offers the most favorable balance of sensitivity, depth, quantitative accuracy, and enrichment fidelity for proximity-labeling proteomics in vivo.

Thus far, studies have typically used 500 µg to 1 mg of protein for each pulldown ^9, 11, 12, 14^. Based on increased peptide recovery in our workflows, we performed experiments to determine the lower limit of starting material required for robust identification of proteins using the CIBOP approach. We applied the optimized SDS on-bead workflow to 100 µg, 80 µg, and 20 µg of input lysate (n = 4: 2 biological and 2 technical replicates per condition) and compared these datasets to our standard full-scale experiment. DEPs showed a high degree of concordance across all input amounts. A core set of 699 proteins was consistently enriched across all conditions, indicating that the most abundant biotinylated targets are reliably captured regardless of input scale (Fig. 3H).

Even at the lowest input of 20 µg, the workflow preserved substantial proteomic depth. The 20 µg samples shared 1,588 proteins with the 100 µg condition, demonstrating strong retention of biological signal. Applying a two-fold enrichment threshold (FC > 2), we identified 3,098 enriched proteins for the 100 µg input and 2,012 enriched proteins for the 20 µg input, with 1,969 proteins overlapping between the two. The reduction in total identifications in the 20 µg cohort was primarily attributable to differences in peptide loading. While the 100 µg digest supported the standard 300 ng analytical injection, the 20 µg digest yielded ∼310 ng total peptide mass. After allocating 10% for peptide BCA quantification and 10% for an MS test injection, <300 ng remained for the analytical run, reducing column load relative to the full-scale workflow.

Despite this lower load, the 20 µg samples maintained strong proteomic depth and reproducibility (Supp Table: 2). These results demonstrate that CIBOP can be successfully performed with as little as 20 µg of starting material, enabling the characterization of proteomes from limited cell populations, micro-dissected tissue samples, or other contexts where material is inherently limited.

### Deep neuronal and synaptic proteomes recovered using optimized TurboID SDS S-Trap DIA workflow

To evaluate the performance of the optimized SDS–S-Trap–DIA workflow to another brain cell type, we applied CIBOP to existing brain tissues from Camk2a-CreERT2/TurboID mice, enabling proximity labeling specifically in forebrain excitatory neurons. Given the high compartmental specialization of excitatory neurons, we processed two complementary sample types of whole-brain homogenates, capturing global neuronal proteomes, and P2 synaptosomal fractions, pinched-off synaptic terminals containing synaptic vesicles and mitochondria. These samples were originally generated as part of our previously published Camk2a–CIBOP study ^53^, in which 500 µg/sample of cortical proteins were used for streptavidin enrichment. For the present benchmarking, we utilized only 15% of the original bead volume (corresponding to ∼75 µg input) from matched homogenate and P2 fractions to directly compare performance against the prior 500 µg in-solution digestion DDA workflow. Importantly, the use of only 75 µg input (compared to 500 µg required in the earlier in-solution digestion DDA study), allowed us to directly test the low-input performance of the optimized DIA workflow (Supp Table. 3). Western blot (WB) and silver stain analysis of streptavidin-enriched Camk2a whole-neuron samples (homogenate pulldowns) and Camk2a synaptosome samples (P2 fraction pulldowns) showed patterns of strong protein biotinylation in Camk2a-CIBOP, with minimal signal in Cre-negative controls (Fig. 4A), providing a strong foundation for assessing neuronal proteome recovery.

**Figure 4:**
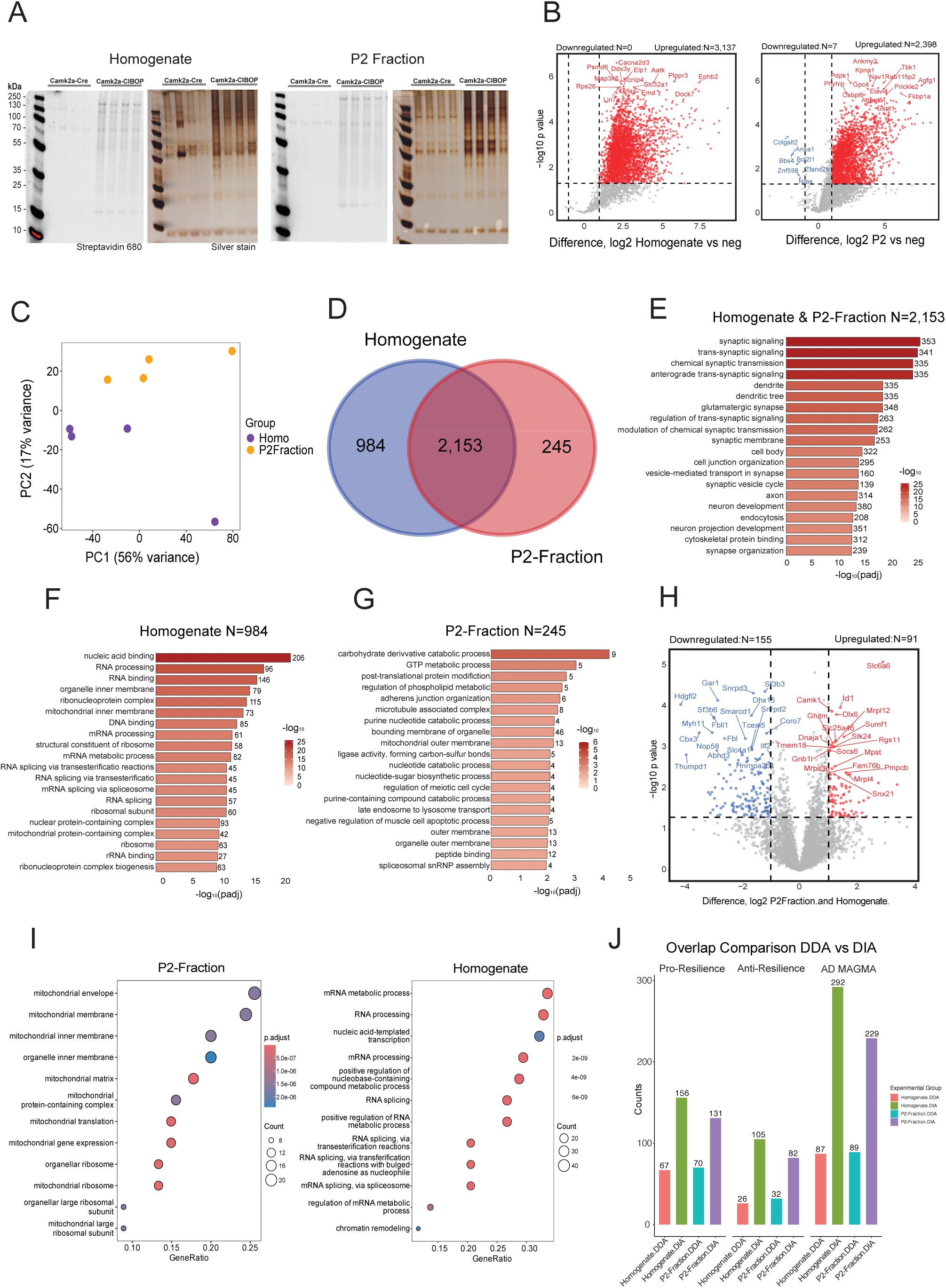
Compartment-resolved neuronal proteomics with Camk2a-TurboID and DIA reveals synaptic and mitochondrial enrichment: **(A) Western blot analysis (left panels) and silver stain (right panels)** of streptavidin-enriched samples from brain homogenates and P2 synaptosomal fractions of Camk2a-CreERT2/TurboID mice (Camk2a-CIBOP) and Cre-negative controls (Camk2a-Cre). Immunoblotting with streptavidin-680 confirms robust and Cre- dependent biotinylation in both homogenate and P2 fractions, demonstrating efficient neuronal proximity labeling across compartments. **(B) Volcano plots depicting differential protein enrichment in Camk2a-CIBOP samples** relative to Cre-negative controls for whole homogenate (left) and P2 fraction (right). Significantly enriched proteins are highlighted in red (fold change ≥2, *p* ≤0.05). The optimized DIA workflow identified **3,137 proteins** in homogenate and **2,398 proteins** in P2 fractions. **(C) Principal component analysis (PCA) of neuronal proteomes** showing dominant separation by cellular compartment along PC1 (56% variance) and secondary separation by experimental group along PC2 (17% variance), indicating strong compartment-specific proteomic signatures. **(D) Venn diagram illustrating overlap between proteins identified in homogenate and P2 fractions.** A shared core neuronal proteome of **2,153 proteins** is detected across both compartments, alongside compartment- specific protein pools (**984 homogenate-enriched** and **245 P2-enriched proteins**). **(E) Gene Ontology enrichment analysis of the shared neuronal proteome (**N = 2,153): reveals enrichment of synaptic organization, vesicle-mediated transport, and trans-synaptic signaling pathways. **(F) Gene Ontology enrichment analysis of homogenate specific proteome (N=984):** RNA processing, and ribonucleoprotein complex assembly. **(G) Gene Ontology enrichment analysis of P2-fraction specific proteome (N=245):** post-transcriptional regulation in homogenate, GTP metabolic processes, organellar membranes, and structural organization. Bar length represents Z-score, and color scale indicates −log10(FDR). **(H) Volcano plot of the direct comparison between P2-fraction and Homogenate samples**. Red dots indicate proteins significantly enriched in P2-fraction (91), and blue dots indicate proteins enriched in Homogenate (155). P2-enriched proteins are associated with mitochondrial and organellar membranes, while Homogenate-enriched proteins are associated with RNA processing and splicing. (**I) Gene ontology enrichment analysis of P2-fractioned and homogenate enriched proteins**. The P2-fraction showed strong enrichment on mitochondrial membranes, mitochondrial matrix, and mitochondrial ribosome. The homogenate enrichment largely was enriched in mRNA metabolic process, RNA processing, RNA splicing. **(J) Comparison of neuronal proteins annotated for Alzheimer’s disease resilience** (Pro-resilience and Anti- resilience) and genetic risk (AD MAGMA) recovered by optimized DIA versus prior DDA workflows in homogenate and P2 fractions. DIA consistently captures a greater number of disease-relevant proteins across all annotation categories, highlighting improved sensitivity for biologically relevant neuronal pathways.

PCA of proteomes demonstrated that tissue compartment was the primary driver of proteomic variance (PC1, 56%), whereas PC2 (17%) effectively resolved treatment groups within each compartment, indicating that synaptic and somatodendritic fractions harbor distinct baseline molecular signatures (Fig. 4C). Using 8 M urea lysis and S-Trap DIA-MS, the optimized workflow identified a total of 7,612 proteins across homogenate and P2 fractions from Camk2a neurons, with specific differentially expressed proteins (DEPs) (FC ≥2, (p ≤ 0.05)) over Cre-negative controls. We quantified 3,137 DEPs in homogenate and 2,398 DEPs in P2 fractions with very few proteins enriched in the Cre-only negative control tissues (Fig. 4B). In comparison, our previous in-solution DDA workflow with 500 µg input detected 860 differentially enriched proteins in P2 saline, 872 in homogenate saline (Supp Fig 4A) ^53^. Thus, the optimized DIA-based method increased proteome coverage by nearly 4-fold and reduced input protein needed by 7- fold.

Comparing homogenate and P2 cellular compartments revealed compartment-specific proteomic signatures. We identified 2,153 proteins detected in both homogenate and P2, along with 984 proteins enriched in homogenates and 245 proteins only enriched in P2 fractions (Fig. 4D). GO analysis of the shared compartment proteins showed strong enrichment for synaptic signaling, trans-synaptic signaling, chemical synaptic transmission, dendrites, glutamatergic synapses, synaptic membranes, axon, synaptic vesicle cycle, and vesicle-mediated transport in synapses (Fig. 4E).Proteins enriched in the homogenate fraction, were dominated by nucleic acid binding, RNA processing and splicing, ribonucleoprotein complexes, ribosomal components, and nuclear and mitochondrial protein-containing complexes (Fig. 4F), reflecting the strong representation of somatic gene expression and translational machinery. In contrast, P2-enriched proteins were associated with carbohydrate derivative catabolic processes, GTP metabolic process, post-translational protein modification, regulation of phospholipid metabolism, adherens junction organization, microtubule-associated complexes, and organelle outer membranes, including mitochondria, along with late endosome to lysosome transport and peptide binding (Fig. 4G). Together, these findings show that while major synaptic pathways are shared, the homogenate compartment emphasizes somatic nuclear and translational processes, whereas the P2 fractions representing the synaptic compartment, enhance detection of membrane- and organelle-associated functions that support structural organization and trafficking at excitatory synapses.

We next performed a direct differential comparison between P2 synaptosomal fractions and whole homogenates to define compartment-specific protein enrichment. With our DEP cutoff, the analysis identified 91 proteins enriched in P2 fractions and 155 proteins enriched in homogenates, consistent with distinct subcellular specialization between the two preparations (Fig. 4H). GO enrichment revealed that P2-enriched proteins were predominantly associated with mitochondrial envelope and membrane organization, mitochondrial ribosomes, and amino acid and protein metabolism, reflecting the high metabolic demand of synaptic compartments ^54, 55, 56^ ^57^. These results reinforce the enhanced ability of P2 enrichment to isolate synaptic and membrane-resolved neuronal components that are not readily detected in whole tissue.

To determine whether Camk2a-labeled neuronal proteins align with genetic determinants of Alzheimer’s disease susceptibility and resilience ^52^, we quantified their overlap with AD MAGMA genes and resilience-associated signatures ^51, 58^. Across all categories, the updated DIA workflow recovered substantially more relevant proteins than the earlier DDA dataset. In homogenates, pro-resilience overlap increased from 67 to 156, anti-resilience from 26 to 105, and AD MAGMA from 87 to 292 ^52, 59^. P2 fractions showed a similar pattern, with increases from 70 to 131 (pro-resilience), 32 to 82 (anti-resilience), and 89 to 229 (AD MAGMA) (Fig. 4J). P2 samples also showed higher absolute overlap than homogenates, indicating that synaptic and membrane-enriched neuronal compartments contain a concentrated set of AD-linked and resilience-associated proteins. These findings demonstrate that Camk2a TurboID robustly captures neuron-intrinsic molecular programs relevant to AD risk and resilience, and that DIA profiling improves recovery of these biologically meaningful insights for cell-specific functions.

## DISCUSSION

Proximity-dependent biotinylation methods for native-state proteomics have emerged as a powerful strategy for resolving cell-type– and compartment-specific proteomes within complex tissues, yet the practical implementation remains limited by longstanding technical barriers related to the tissue lysis, enrichment, digestion, and data acquisition ^9,8^. In this study, we performed a rigorous and comprehensive, side-by-side optimization of the CIBOP workflow, spanning in vitro microglial models, astrocyte and neuronal specific in vivo labeling, by cytosolic TurboID. By systematically comparing lysis chemistries, digestion strategies, affinity purification formats, mass spectrometry acquisition modes, and instrument architectures, we define an integrated pipeline that addresses these bottlenecks and substantially expands the accessible proteomic landscape of TurboID-labeled cells.

Our careful benchmarking studies also establish on-and off-beads S-Trap digestion as the most reliable and reproducible approaches for processing TurboID-based proximity labeling samples. On-bead digestion minimizes sample transfers, reduces stochastic loss of hydrophobic peptides, and increases digestion efficiency, which together yields improved quantitative precision relative to off-bead workflows. Importantly, the concern of streptavidin peptide contamination introduced by on-bead digestion is functionally irrelevant in DIA-MS, where comprehensive fragmentation and signal deconvolution suppress interference. The ∼20-fold increase in peptide yield compared with in-solution digestion further reinforces S-Trap as a key enabling technology for proximity proteomics, especially when starting material is limited.

A central insight from our work is that detergent-based lysis is essential for comprehensive CIBOP profiling in intact brain tissue. While urea lysis was sufficient and only slightly superior to SDS-based lysis to recover most biotinylated proteins in cultured BV2 microglia, it proved inferior to SDS-based lysis for astrocyte proteomic profiling in vivo, where a substantial fraction of the proteome resides within membrane-rich processes, vesicular compartments, perivascular endfeet and perisynaptic endfeet ^32, 33^. SDS lysis overcame these limitations by solubilizing dense synaptic and membrane-proximal structures that 8 M urea lysis cannot efficiently access. This further emphasizes that optimal lysis depends on cellular architecture, with complex neural tissues requiring stronger detergents to extract reveal their full proximity-labeled proteomes. The resulting datasets also provide deep access to astrocyte–neuron interface biology, revealing synaptic scaffolding proteins, signaling adaptors, vesicle-trafficking machinery, and structural regulators that were largely inaccessible using classical urea-based protocols. The enrichment of AD vulnerability, and cognitive resilience, within these SDS-enriched astrocyte datasets further highlights the biological value of this expanded coverage.

At the level of data acquisition, our results provide a clear framework for choosing between DDA, TMT, and DIA strategies. The deterministic sampling of DIA is particularly advantageous for the highly skewed peptide populations produced by TurboID enrichment, avoiding the stochasticity that hampers DDA and ratio of compression that undermines TMT. The additional gains achieved with the Orbitrap Astral consistently identifying hundreds to thousands more enriched proteins across conditions highlight how instrument architecture directly influences the depth and interpretive potential of proximity labeling studies. Together, these observations support the conclusion that SDS lysis along with S-Trap digestion and DIA represents the most powerful combination currently available for TurboID-MS proteomics.

A practical strength of the optimized workflow is its robustness at low starting input, enabling high-quality proximity proteomics from as little as 20 µg of lysate. This substantially lowers the barrier for studying rare cell populations, fine neuroanatomical structures, or sparse Cre-driver lines, and opens the possibility of analyzing microdissected tissue regions or small numbers of FACS-isolated cells^14^. Achieving this depth with low input also facilitates experimental designs requiring multiple perturbation groups or longitudinal sampling, which are prohibitive with traditional high-input protocols.

The optimized workflow also enhances biological discovery. In astrocytes, SDS-enabled solubilization exposed perisynaptic and membrane-associated proteomes, allowing us to capture structural, signaling, and regulatory proteins that define astrocyte–neuron crosstalk ^11, 30^. Analysis of P2 synaptosomal fractions and whole homogenates, each processed from only 75 µg of input, demonstrated that the optimized workflow sensitively recovers the major molecular features that define excitatory neurons. Importantly, it preserves clear compartment specificity highlighting synaptic organization and signaling in P2 fractions and broader somatic programs in homogenates—while providing substantially deeper neuronal proteome coverage than prior high-input methods. These molecular signatures align with recent consensus frameworks that define astrocyte reactivity as a multidimensional spectrum rather than a binary state ^31, 60^. Together, the optimized workflow enables deep, reproducible, and compartment-resolved profiling of cells and cellular compartments from minimal input material, substantially outperforming earlier DDA approaches. The detection of synaptic and inflammatory remodeling further establishes its sensitivity for studying physiological perturbations and neuron-specific molecular adaptations.

While our workflow substantially improves sensitivity, membrane coverage, and reproducibility, limitations remain. TurboID labeling radius and expression levels constrain spatial resolution, and SDS-based solubilization precludes the recovery of certain native complexes or biotinylation-sensitive protein states. Future work may integrate orthogonal approaches such as organelle-targeted ligases or chemoproteomic enrichment of PTMs to further refine cell-type and subcellular specificity ^61, 62^. For example, resolving the proteomes of discrete domains like the axon initial segment would provide critical insights into neuronal polarity and excitability that global profiling cannot capture ^63, 64^, while technologies like APEX2 ^65^ and Split-TurboID ^66^ offer enhanced temporal and contact-dependent resolution. The combination of TurboID with ultra-low-input DIA methods may soon enable single-nucleus or sparse-population proximity proteomics, building on the low input compatibility demonstrated here.

In conclusion, this study provides a comprehensive and experimentally validated roadmap for optimizing TurboID-based proximity proteomics across cell culture systems and intact neural tissues. By addressing the key bottlenecks, we establish a workflow that markedly enhances proteome depth and expands the accessible landscape of cell-type–specific proteomes from limited material. This optimized CIBOP platform will support a new generation of studies aimed at unraveling cell-type–specific signaling, metabolic regulation, and molecular vulnerability in health and disease.

## METHODS

### Cell Culture and BV2 TurboID Labeling

BV2 microglial cells were maintained in Dulbecco’s Modified Eagle Medium Nutrient Mixture F-12 (DMEM:F12) (Gilbco,11330-032), L-glutamine, 10% fetal bovine serum (FBS) (Gibco, 26140-079), and 1% penicillin–streptomycin (Gibco,15140-122). All media were sterilized via 0.2 µm vacuum filtration. Cells were cultured at 37 °C in a humidified incubator with 5% CO and passaged at approximately 80% confluency. For mass spectrometry experiments, BV2 cells were expanded in 150 mm dishes to ∼95% confluency. Stable lines expressing cytosolic V5- TurboID-NES were maintained under standard conditions and incubated in medium supplemented with 200 µM biotin for 24 h prior to harvesting. Two biological replicates (n = 2) were processed in parallel for each experimental workflow. Cells were washed twice with ice-cold phosphate-buffered saline (PBS), detached using a cell scraper, pelleted at 15,000 × g for 5 min at 4 °C, and transferred to Eppendorf LoBind tubes for all downstream processing.

### Animals and In Vivo TurboID Labeling

All animal procedures were performed in accordance with Yale University IACUC protocols (PROTO2023-20536) and aligned with NIH guidelines for laboratory animal welfare. Mice were housed in a controlled vivarium environment (72 °F, 40–50% humidity) under a 12-h light/dark cycle with ad libitum access to food and water. Aldh1l1-CreERT2;Rosa26-TurboID and Camk2a-CreERT2;Rosa26-TurboID mice were generated by crossing Aldh1l1-CreERT2 or Camk2a-CreERT2 (JAX #012362) drivers with Rosa26-TurboID (C57BL/6- Gt(ROSA)26Sortm1(birA)Srgj/J, JAX #037890) Tamoxifen (75 mg/kg) was administered via intraperitoneal injection for 5 consecutive days at 8 weeks of age, followed by administration of biotin-supplemented drinking water (37.5 mg/L) for 2 weeks prior to euthanasia. For Aldh1l1- TurboID experiments, mice (n = 4 total; 2 biological replicates × 2 technical replicates) were anesthetized with ketamine/xylazine (87.5 mg/kg ketamine, 12.5 mg/kg xylazine) and transcardially perfused with 30 mL of ice-cold PBS. Brains were removed, hemisected along the mid-sagittal plane, and snap-frozen on dry ice. For Camk2a-TurboID experiments (n = 4 total samples/genotype), mice were deeply anesthetized and decapitated, and their brains were quickly removed, brain homogenates and crude synaptosomes (P2 fraction) were prepared by differential centrifugation as previously described^53^. Briefly, each brain was gently homogenized with 16- strokes at 800 rpm in 10 volumes of Medium I (0.32 M sucrose, 5 mM HEPES pH 7.5, and 0.1 mM EDTA) containing protease and phosphatase inhibitors (ThermoFisher Scientific, 78446). The homogenate was first centrifuged at 1,000 x g for 10 min to give a pellet (P1) containing nuclear and cell debris. Supernatant (S1) was then centrifuged at 12,000 x g for 20 min to produce the crude synaptosome pellet (P2). Supernatant (S2) was discarded, the P2 pellet was resuspended in 500 µL Medium I and sonicated on ice until completely dissolved, and the resulting P2 fractions were stored at -80°C until used for downstream analysis.

### Protein Extraction and Lysis Conditions

Cell pellets or brain tissue samples were lysed under one of three conditions depending on the experimental workflow: (i) Urea lysis using 8 M urea, 10 mM Tris, and 100 mM NaH PO (pH 8.5) supplemented with HALT protease inhibitor (EDTA-free); (ii) SDS lysis using 5% SDS in 50 mM triethylammonium bicarbonate (TEAB); or (iii) sequential urea/SDS lysis, beginning with 8 M urea buffer followed by the addition of SDS buffer to a final concentration equivalent to 5% SDS. Homogenization was performed in LoBind microcentrifuge tubes followed by three cycles of sonication (5 s at 25% amplitude, 10 s rest on ice). Lysates were clarified by centrifugation at 15,000 × g for 5 min at 4 °C, and protein concentration was quantified using the Pierce BCA assay. Total protein aliquots of 500 µg (cell lysates) or 1,000 µg (brain lysates) were prepared for each experimental condition and stored at −80 °C until enrichment.

### Streptavidin Enrichment of Biotinylated Proteins

Streptavidin magnetic beads (Thermo 88817) were pre-washed twice in RIPA buffer (50 mM Tris pH 7.4, 150 mM NaCl, 0.1% SDS, 0.5% sodium deoxycholate, 1% Triton X-100). Beads were added to lysates (500/1,000 µg) at a protein: bead ratio of 11.8:1 (42.5 µL bead slurry per sample) and incubated for 1.5 hour at 4 °C with rotation. Samples derived from SDS lysis were diluted with RIPA buffer lacking SDS to a final SDS concentration of 0.2% prior to enrichment. Following incubation, supernatants were removed, and beads were subjected to a sequential high-stringency wash protocol at room temperature consisting of: two washes with RIPA buffer (8 min each), one wash with 1 M KCl (8 min), one brief (1 min) wash with 0.1 M Na CO , one wash with (1 min) 2 M urea in 10 mM Tris-HCl (pH 8.0), two additional 8-min washes with RIPA buffer, two washes with PBS (2 min each), and five manual PBS rinses. The final bead suspension was normalized to 100 µL in PBS and divided according to the designated workflow. For S-Trap experiments, the suspension was split into two 45 µL aliquots for parallel on-bead and off-bead digestion. For in-solution digestion (ISD) experiments, 90 µL of the suspension was utilized. In all workflows, the remaining 10 µL aliquot was eluted with 2× SDS sample buffer containing 2 mM biotin and 10 mM DTT at 95 °C for 10 min to assess enrichment quality via streptavidin blotting and silver staining.

### S-Trap and In-Solution Digestion and Quantification

For S-Trap workflows, beads were magnetically separated and resuspended in 5% SDS in 50 mM TEAB. Proteins were reduced by 10 mM DTT (1 h, room temperature) and alkylated with 20 mM iodoacetamide (IAA) (30 min, dark). For off-bead digestion, proteins were eluted in 2 mM biotin with 10 mM DTT at 95 °C for 5 min prior to addition of IAA. Samples were acidified to 1.2% phosphoric acid, mixed with binding buffer (90% methanol, 100 mM TEAB, pH 7.1), loaded onto S-Trap microcolumns in 70 µL increment till full transfer (Protifi) (19 Zougman), and washed three times. Digestion was performed sequentially with Lys-C (0.5 µg, 4.5 h at 37 °C) and trypsin (1 µg, 18 h at 37 °C), followed by peptide elution. For ISD workflows, 90% of streptavidin beads were reduced, alkylated, diluted fourfold with 50 mM ammonium bicarbonate, and digested with Lys-C and trypsin as described above. Peptides were desalted using HLB cartridges, dried, and reconstituted for analysis. Dried peptides were resuspended in 1% formic acid in HPLC-grade water, and peptide concentration was determined using the Pierce Quantitative Fluorometric Peptide Assay (Thermo Scientific, 23290) according to the manufacturer’s protocol.

### TMT labeling

After drying the sample, the peptides were isotopically labeled using a 10-plex tandem mass tag (TMT, Thermo, 90110)^22^.The reaction proceeded for 2 hours at room temperature at an intermediate vortex speed, and the sample was spun down every 15 min. The labeled samples were pooled, resuspended in 50 mM TEAB buffer, and then fractionated using the Thermo Scientific Pierce High pH Reversed-Phase Peptide Fractionation Kit (Thermo Scientific, 84868).

### Liquid Chromatography Configuration

LC–MS/MS analyses were performed on a Vanquish Neo UHPLC system coupled to an Orbitrap Exploris 480 mass spectrometer (Thermo Fisher Scientific) equipped with an EASY-Spray source. To ensure column performance, a fixed injection volume was determined based on the sample with the highest peptide concentration, ensuring a maximum total peptide load of 300 ng per injection. Samples were loaded onto a PepMap Neo trap column (300 μm i.d. × 5 mm, 5 μm C18) and separated on an Aurora Frontier analytical column (75 μm i.d. × 60 cm, 1.7 μm C18, IonOpticks) at a flow rate of 300 nL/min. Mobile phases consisted of 0.1% formic acid in water (Phase A) and 0.1% formic acid in 80% acetonitrile (Phase B). Peptides were eluted using the following gradient: 1–4% B (10 min), 4–25% B (60 min), and 25–65% B (45 min), followed by a column wash (65–99% B in 1 min, held for 19 min) and re-equilibration. The ESI voltage was set to 2,100 V, and the ion transfer tube temperature was 275 °C. Internal calibration was performed using the lock mass of polydimethylcyclosiloxane (m/z 445.12003).

### DIA Acquisition Parameters

MS analysis was performed in data-independent acquisition (DIA) mode. The method consisted of one MS1 scan (m/z 400–1000) followed by 60 DIA MS2 scans. MS1 spectra were acquired at a resolution of 120,000 (at m/z 200) with a normalized AGC target of 300% (3 × 10 charges), an ‘Auto’ maximum injection time, and an RF lens setting of 50%. The 60 DIA windows were acquired at a resolution of 30,000 with a normalized AGC target of 1000% (1 × 10 charges) and ‘Auto’ maximum injection time. Precursor ions were isolated using a 10 m/z window with 1 m/z overlap and fragmented via high-energy collision dissociation (HCD) at a normalized collision energy (NCE) of 28%.

### DDA Acquisition Parameters

For data-dependent acquisition (DDA), the instrument operated with automatic switching between MS1 and MS2 scans. MS1 spectra (m/z 350–1500) were acquired at a resolution of 60,000 with a normalized AGC target of 300% (3 × 10 charges), a maximum injection time of 50 ms, and an RF lens setting of 50%. The top 20 most intense precursors (charge states 2–7; minimum intensity 2 × 10 ) were selected for isolation (0.7 m/z window) and fragmentation (HCD, NCE 32%). MS2 spectra were acquired at a resolution of 45,000 with a normalized AGC target of 50% (5 × 10 charges) and a maximum injection time of 86 ms. The first mass was fixed at m/z 120. Dynamic exclusion was set to 25 s to minimize repeated sequencing of abundant peptides.

### Data Processing using DIA-NN

The acquired DIA raw files were processed using DIA-NN (v2.2.0, Academia) in library-free mode (23 Demichev). The search was conducted against a Mus musculus (Mouse) FASTA database (UniProt UP000000589, downloaded 2025.04.11) supplemented with common contaminants. Search parameters were set to Trypsin/P specificity, allowing for up to two missed cleavages. Carbamidomethylation (C) was set as a fixed modification. Variable modifications included N-terminal M excision, Oxidation (M), and N-terminal acetylation, with a maximum of three variable modifications per peptide. The peptide length was restricted to 7–30 amino acids, and the precursor charge range was set from 1 to 4. The analysis matched precursors within the m/z 400–1000 range and fragments within the m/z 150–2000 range. Mass tolerances for both MS1 and MS2 were set to 10 ppm, with a scan window of 15. The analysis used neural network (NN)-based, cross-validated machine learning and proteoform-level scoring. All results were filtered at a 1% precursor and protein group of FDR. Protein quantification was performed using the QuantUMS strategy. Cross-run normalization was disabled within DIA-NN.

### Proteomic discoverer and CHIMERYS

Raw data were processed in Proteome Discoverer 3.2 (Thermo Fisher Scientific) (47 Thermo Fisher Scientific) using CHIMERYS-based workflows optimized for data-dependent acquisition (DDA). Spectra were searched with CHIMERYS v4.0.24 using the inferys 4.7.0 fragmentation prediction model against the UniProt Mus musculus canonical proteome (release 408, 18 June 2025) supplemented with a Thermo contaminant database (PD_Contaminants_KBH_013125.fasta). Trypsin/P was specified as the digestion enzyme with full specificity, allowing up to two missed cleavages. Precursors and fragment mass tolerances were set to 20 ppm. Carbamidomethylation of cysteine was included as a static modification and methionine oxidation as a variable modification (maximum three variable modifications per peptide). PSMs and peptides were filtered using a target–decoy strategy and controlled at a 1% strict false discovery rate. For TMT-labeled samples, static modifications also included TMT10plex tags on peptide N-termini and lysine residues. Reporter ion intensities were extracted using the Reporter Ion Quantifier node with a 20 ppm integration tolerance, HCD-only MS2 selection, and the “most confident centroid” integration method. Peptide abundance normalization and co-isolation interference correction were performed automatically within Proteome Discoverer. For label-free datasets, Minora feature tracing was used to extract MS1 peptide abundance features (minimum trace length 5 scans, signal-to-noise ≥2, PSM confidence ≥High).

### Protein filtering, imputation, and differential enrichment analysis

For each experimental condition, protein abundance matrices generated by BV2 Astral method analysis have been investigated in R program. An initial filtering step based on data completeness in the positive Turbo samples was implemented to lessen the impact of sparsely identified proteins. Since these characteristics were deemed unreliable for determining Turbo- specific enrichment, proteins with missing values in more than 50% of Turbo replicates were eliminated. To preserve a uniform collection of measured proteins for comparison, the resulting filtered protein list was then applied consistently to all samples, including BV2 controls. After filtering, a Perseus-style binomial imputation method intended to simulate low-abundance signal loss was used to impute missing values. This approach assumed that missingness mostly results from proteins falling below the detection limit rather than actual absence, hence missing intensities were substituted with values taken from a left-shifted distribution in relation to the observed data. This imputation technique maintained the general structure of the abundance distributions while enabling downstream fold-change computations. Group labels were harmonized across experiments, and mean protein abundances were calculated separately for Turbo and BV2 replicates. Differential enrichment was evaluated using log2 fold change, defined as the difference between Turbo and BV2 mean abundances. Proteins with log2 fold change ≥ 1 were considered upregulated, those with log2 fold change ≤ -1 were considered downregulated, and all others were classified as unchanged. For each experiment, protein-level summaries including group means, fold changes, and classification labels were exported for downstream analysis. To compare enrichment patterns across experiments, the number of upregulated proteins was summarized and visualized using bar plots, and results were compared with an independent Exploris480 two-fold enrichment analysis to assess concordance between pipelines. We used differential up-regulated proteins at a 2-fold change from each experiment for overlap with pro-resilience, anti-resilience, and AD Magma list for detection of neurodegenerative proteins.

We applied the same method in Aldh1 TMT and in low amount method samples (20, 80 and 100) to analyze the replicates-based fold changes.

### DIA: filtering, imputation and DEP analysis for astrocytic tissue samples

Proteins were filtered using Turbo samples only to remove features not reliably detected in the positive condition. When a protein abundance value was greater than zero, it was deemed present. If a protein was found in at least 50% of Turbo replicates, it was retained. This list of retained proteins was then applied to the entire dataset, which included both Turbo and negative samples. To simulate signal drop-out close to the detection limit, filtered matrices were imputed using a Perseus-style binomial imputation technique, substituting low-intensity values for missing values. The imputed matrices were used for all downstream statistics. The differential abundances protein one-way ANOVA approach was used to identify differentially expressed proteins. The Benjamini-Hochberg method was used to compensate for multiple testing. When a protein satisfied both requirements nominal p < 0.05 and |log2FC| > 1 (Turbo vs. negative). It was deemed significant. For all experiments, volcano plots and outcome tables were produced.

### Detection of percent of CV

Only Turbo samples were used to measure repeat variability for Aldh1 DIA data. For each protein, the mean and standard deviation across Turbo replicates were computed utilizing imputed, Turbo-only protein abundance tables. The coefficient of variation (%CV) was calculated as (SD/mean) × 100. Violin plots were employed to integrate all per-protein %CV results from all experiments and evaluate replicate consistency across varied experimental conditions.

### Filtering, imputation and DEP analysis for neuronal tissue samples

Camk2a proteomics data were analyzed across all samples (Camk2a-positive and Cre-negative). Proteins were first filtered based on detection consistency in the Camk2a-positive group, retaining proteins quantified in at least 50% of positive samples; this filtered feature set was subsequently used for all analyses. Within each condition, protein intensities were log-transformed, and missing values were imputed in perseus using an identical workflow applied to both positive and negative samples. Following imputation, datasets were subdivided into Homo and P2 fractions, and median normalization was applied independently within each subset. Differential protein analysis was performed between Camk2a-positive and negative groups using one way ANOVA-based method, with statistical significance defined at p < 0.05 and fold-change thresholds applied as specified. GO enrichment analysis was performed using cluster Profiler with mouse annotation (36 Yu). Enrichment was tested against a predefined background universe across all domains (BP, MF, CC). To retain the full-term space for ranking and visualization, no multiple-testing correction was applied (adjustment = “none”), and permissive cutoffs were used (pvalueCutoff = 1; qvalueCutoff = 1). GO results were exported as a complete table, and terms were ranked using the smallest available p-value (adjust when present, otherwise raw p). Enrichment significance was visualized as -log10(adjusted P-value), with the top enriched terms plotted as horizontal bars, bar length reflecting -log10(adjusted P-value)) and labels indicating the number of proteins contributing to each term (Count).

## Supporting information

Table_Legends_Main_Figures

## Abbreviations

AD: Alzheimer’s disease
Aldh1l1: aldehyde dehydrogenase 1 family member L1
BV2: BV2 mouse microglial cell line
Camk2a: calcium/calmodulin-dependent protein kinase II alpha
GPCR: G protein–coupled receptor
MAGMA: Multi-marker Analysis of GenoMic Annotation
PSD: postsynaptic density
AP: affinity purification
BCA: bicinchoninic acid
CIBOP: Cell-type–specific in vivo Biotinylation of Proteins
DMEM: Dulbecco’s Modified Eagle Medium
DTT: dithiothreitol
ESI: electrospray ionization
FACS: fluorescence-activated cell sorting
FASP: filter-aided sample preparation
FBS: fetal bovine serum
NES: nuclear export signal
PBS: phosphate-buffered saline
RIPA: radioimmunoprecipitation assay
SDS: sodium dodecyl sulfate
S-Trap: Suspension-Trapping
SP3: Single-Pot, Solid-Phase–enhanced Sample Preparation
TEAB: triethylammonium bicarbonate
TMT: Tandem Mass Tag
TurboID: engineered biotin ligase TurboID
AGC: automatic gain control
CV: coefficient of variation
DDA: data-dependent acquisition
DIA: data-independent acquisition
FDR: false discovery rate
GO: Gene Ontology
HCD: higher-energy collisional dissociation
LC–MS/MS: liquid chromatography–tandem mass spectrometry
LFQ: label-free quantification
MS: mass spectrometry
PCA: principal component analysis
UHPLC: ultra-high-performance liquid chromatography

## DECLARATIONS

### Ethics approval and consent to participate

Not applicable.

### Consent for publication

Not applicable.

### Availability of data and materials

The MS proteomic data have been deposited to the ProteomeXchange Consortium and can be accessed via the PRIDE data repository with the dataset identifier, project accession: PXD072833. The Rstudio codes for this analysis will be provided upon request.

### Author Contributions

W.E.J. designed the study, performed experiments, developed and optimized mass spectrometry workflows, analyzed data, generated figures, and wrote the manuscript. U.S. performed computational analyses, analyzed data and contributed interpretation. D.K. and R.K. assisted with experimental execution and data validation. P.K. and C.E.-G. provided key mouse lines and contributed to experimental design. A.D.B. assisted with data visualization and validation. S.R. supervised the project, secured funding, guided scientific direction, and edited the manuscript. All authors discussed the results and approved the final manuscript.

## Acknowledgements

We thank Yale Keck Mass Spectrometry & Proteomics Resource for providing access to computational resources, including CHIMERYS and Proteome Discoverer version 3.2, which supported the DDA and TMT database searches. We thank Dr. Katie Henke for technical assistance with data processing through the core facility. We also thank members of the Rangaraju laboratory for helpful discussions and feedback throughout this project.

