## Supplementary figures and images for "Deeper neuronal and glial proteomic insights using an optimized pipeline for proximity labeling proteomics"

### Supplementary_Figures

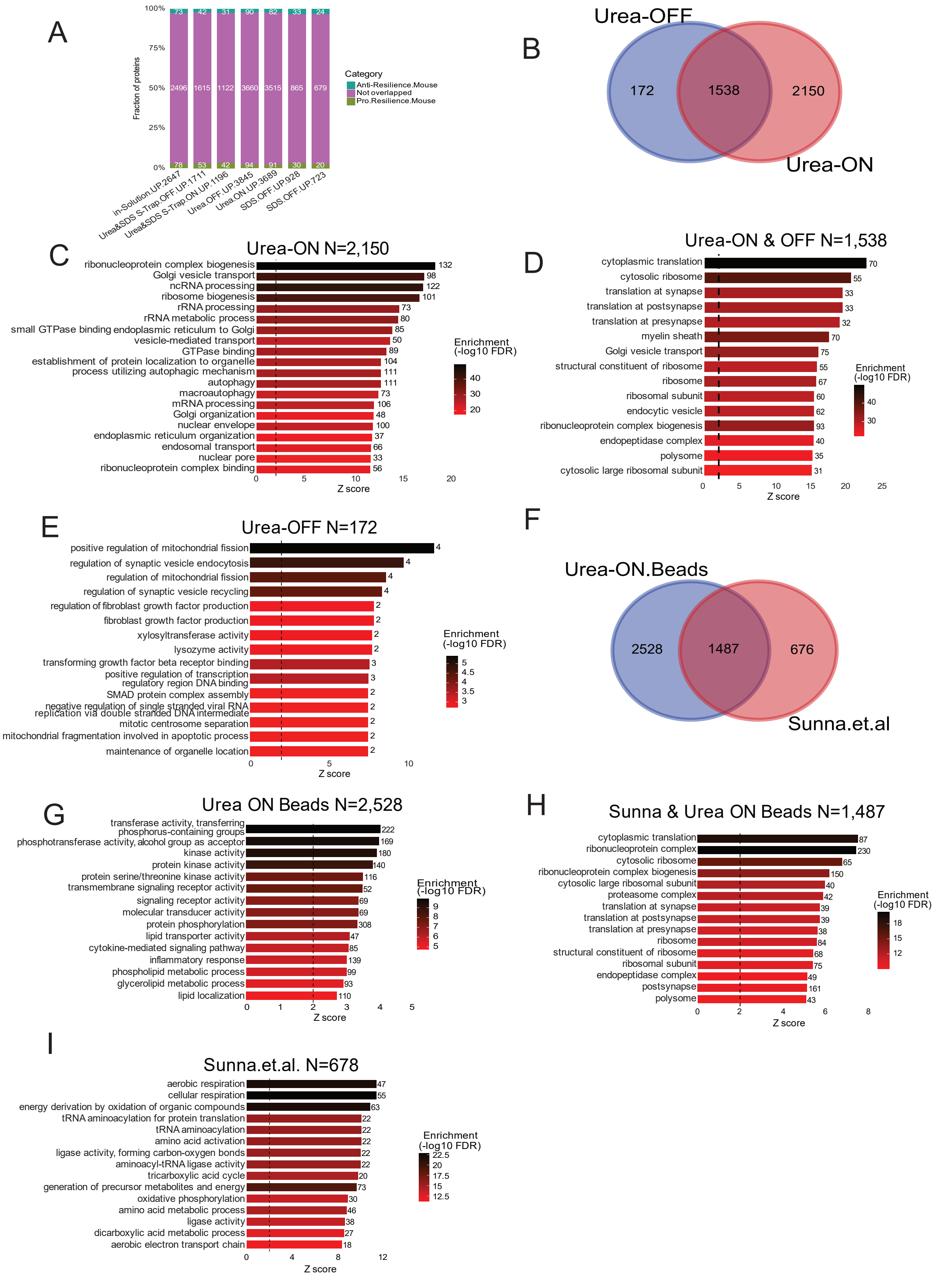

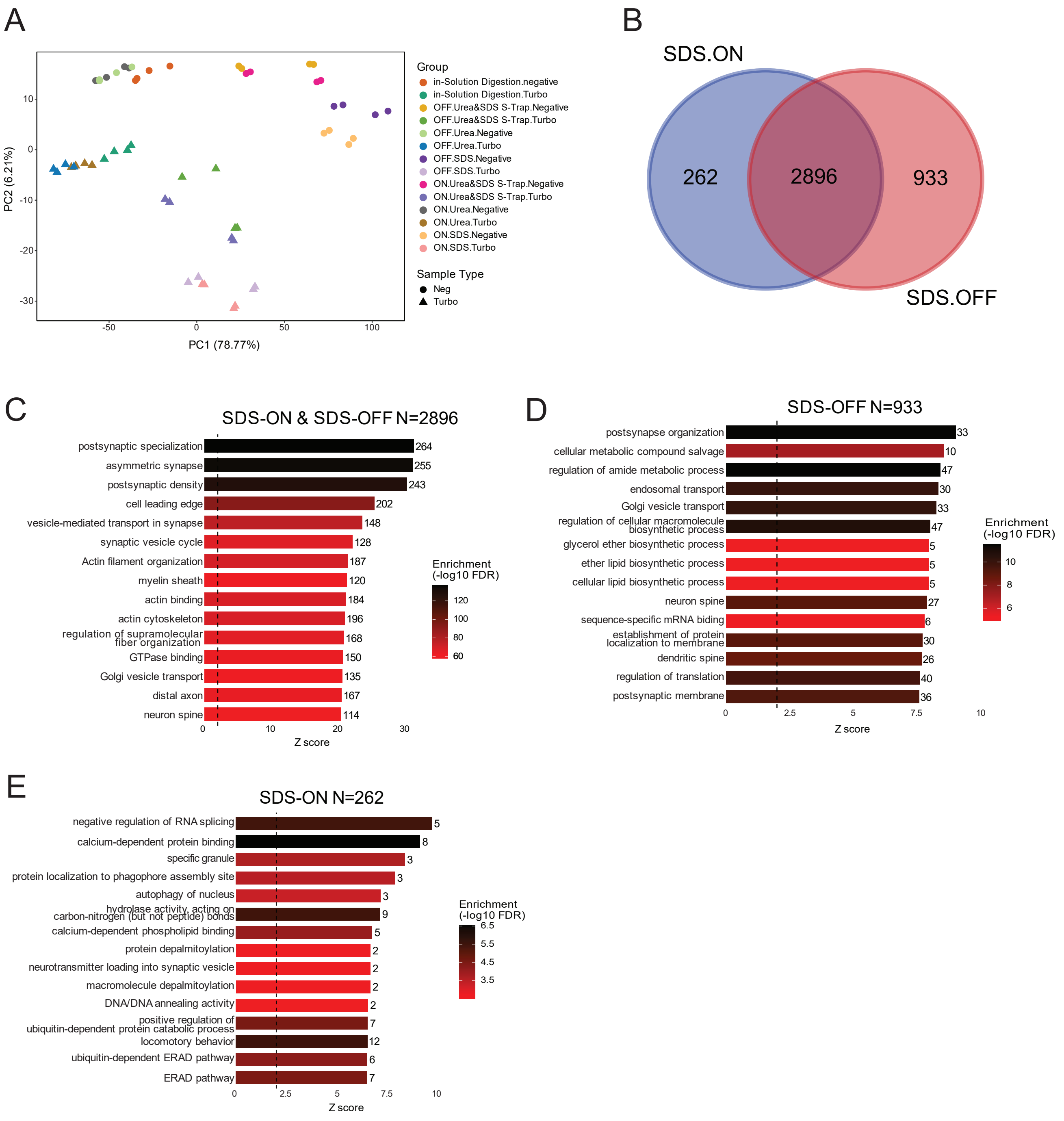

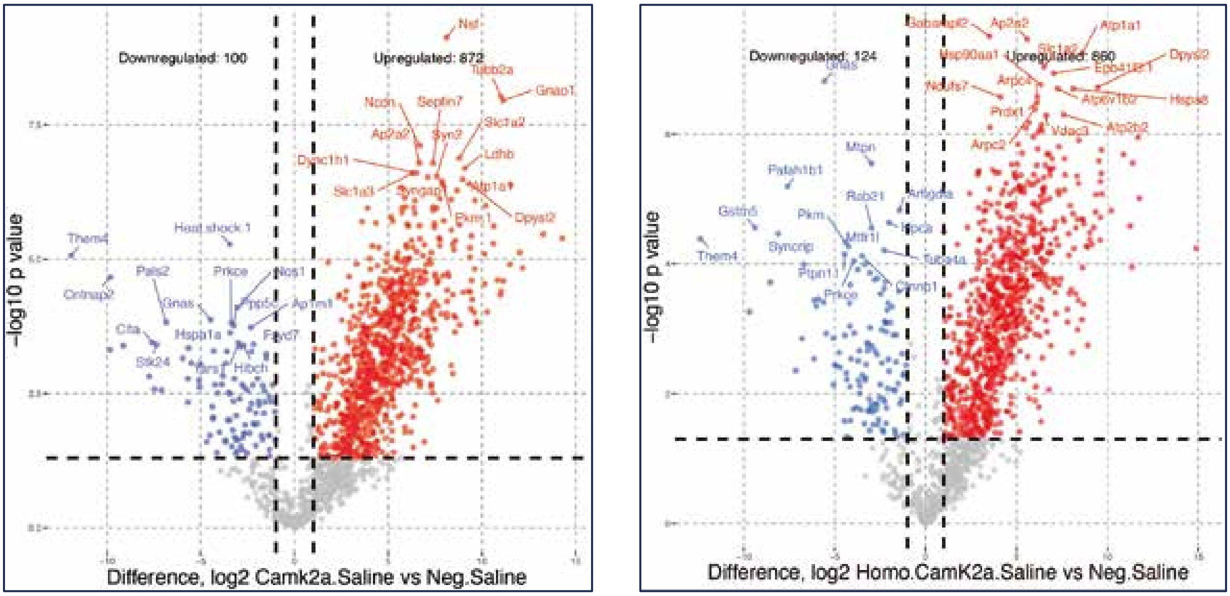
