## Supplementary material for "Deeper neuronal and glial proteomic insights using an optimized pipeline for proximity labeling proteomics": Table_Legends_Main_Figures

Table Legends: Datasheets

Datasheet 1: Data analysis for Figure 1

Sheet #1 (Metadata_BV2): Sample metadata for BV2 TurboID experiments.

Sheet #2 (Peptide_BCA_BV2): Peptide concentration measurements for BV2 samples.

Sheet #3 (raw_BV2_Exploris480): Raw intensity of BV2 Exploris 480 before process

Sheet #4 (raw_BV2_Astral): Processed proteomics data for BV2 samples analyzed on the Astral.

Sheet #5 (Fig. 1C): Principal component analysis (PCA) of BV2 Exploris 480 proteomes across all conditions.

Sheet #6 (Fig. 1D): Bar plot of protein identification counts at baseline (FC = 0) and ≥2-fold change.

Sheet #7 (Fig. 1E): Coefficient of variation distributions across conditions for BV2 samples analyzed on the Exploris 480.

Sheet #8(Fig. 1F): UpSet plot showing protein overlap between conditions using an FC ≥2 cutoff.

Sheet #9 (Fig. 1G): Gene ontology (GO) enrichment analysis of proteins identified following urea lysis.

Sheet #10 (Fig. 1H): GO enrichment analysis of proteins identified following urea lysis with S-Trap digestion.

Sheet #11 (Fig. 1I): GO enrichment analysis of the core proteome (506) shared across all experimental conditions.

Sheet #12 (Fig. 1J): Bar plot for comparison of protein identification counts between Astral and Exploris 480 mass spectrometry platforms.

Datasheet 2: Data analysis for Figure 2

Sheet #1 (Metadata_Aldh1l1): Sample metadata for Aldh1l1-TurboID experiments.

Sheet #2 (Raw_Aldh1l1): Processed proteomics data for Aldh1l1-TurboID samples.

Sheet #3 (Peptide_BCA_Aldh1l1): Peptide concentration measurements for Aldh1l1-TurboID samples.

Sheet #4 (Fig. 2C): Coefficient of variation distributions across experimental conditions in Aldh1l1 samples.

Sheet #5 (Fig. 2D): PCA of Aldh1l1 proteomes across all conditions.

Sheet #6 (Fig. 2E): Bar plot of protein identification counts at baseline (FC = 0) and ≥2-fold change.

Sheet #7 (Fig. 2F): UpSet plot showing protein overlap between conditions using an FC ≥2 cutoff.

Sheet #8 (Fig. 2G): GO enrichment analysis of proteins (1,500) identified with S-trap digestion with SDS lysis

Sheet #9 (Fig. 2H): GO enrichment analysis of proteins (692) identified only identified with S-trap digestion with SDS On-beads

Sheet #10 (Fig. 2I): GO enrichment analysis of proteins (257) identified through all core experiments

Datasheet 3: Data analysis for Figure 3

Sheet#1 (Low_Peptide_BCA): Peptide BCA of 20, 80, and 100 ug affinity pulldown – digestion, peptide BCA.

Sheet #2 (Fig. 3A): Venn diagram comparing SDS–S-Trap–DIA data with previously published differentially expressed proteins identified using urea lysis, in-solution digestion, and DDA (Rayaprolu et al., 2022).

Sheet #3 (Fig. 3B): GO enrichment analysis of proteins (2,280) uniquely identified by SDS–S-Trap–DIA.

Sheet #4 (Fig. 3C): GO enrichment analysis of proteins (878) commonly identified across SDS–S-Trap–DIA and previously published methods.

Sheet #5 (Fig. 3D): GO enrichment analysis of proteins (427) uniquely identified by urea lysis with in-solution digestion and DDA.

Sheet #6 (Fig. 3E): Bar graph differentially enriched proteins in Aldh1l1 across methods highlighting enrichment of neurological disorder–associated resilience genes.

Sheet #7 (Fig. 3F): Comparison of TMT, LFQ, DDA, and DIA workflows demonstrating highest protein recovery with SDS-based DIA.

Sheet #8 (Fig. 3G): Venn diagram comparing TMT, LFQ-DDA, and DIA workflows showing DIA captures the largest unique proteome with a shared core across methods.

Sheet #9 (Fig. 3H): Venn diagram comparing low-input DIA datasets.

Datasheet 4: Data analysis for Figure 4

Sheet #1 (Meta_Camk2a): Sample metadata for Camk2a-TurboID experiments.

Sheet #2 (Peptide_BCA_Camk2a): Peptide concentration measurements for Camk2a-TurboID samples.

Sheet #3 (Camk2a_Proteomic_Data): Raw Intensity Camk2a pull-down

Sheet #5 (Fig. 4B): Volcano plot comparing homogenate and P2 fractions from Camk2a-TurboID samples against Camk2a-Cre controls.

Sheet #6 (Fig. 4C): Camk2a PCA of homogenate and P2 fraction proteomes.

Sheet #7 (Fig. 4D): Venn diagram of differentially expressed proteins between homogenate and P2 fractions.

Sheet #8 (Fig. 4E): GO enrichment analysis of proteins (2,153) shared between homogenate and P2 fractions.

Sheet #9 (Fig. 4F): GO enrichment analysis of proteins (984) uniquely identified in the homogenate proteome.

Sheet #10 (Fig. 4G): GO enrichment analysis of proteins (245) uniquely identified in the P2 fraction proteome.

Sheet #11 (Fig. 4H): Venn diagram comparing protein identifications between homogenate and P2 fractions.

Sheet #12 (Fig. 4I): GO enrichment terms for differentially expressed proteins in homogenate and P2 fractions.

Sheet #13 (Fig. 4J): Bar plot comparing the number of identified proteins annotated for Alzheimer’s disease resilience and genetic risk factors.
