## Supplementary material for "Deeper neuronal and glial proteomic insights using an optimized pipeline for proximity labeling proteomics": Table_Legends_Supplementary_Figures

Datasheet 1: Data analysis for Figure 1 Supplementary

Sheet 1 (Fig. 1A.Supp): Bar plot: overlap of proteins associated with resilience across BV2 experimental conditions.

Sheet 2 (Fig. 1B.Supp): BV2 Exploris480 PCA of distinct clustering by lysis and digestion method.

Sheet 3 (Fig. 1C.Supp): Venn diagram Overlap of proteins identified in Urea ON- and OFF-bead fractions.

Sheet 4 (Fig. 1D.Supp): GO enrichment analysis of urea ON-bead–specific proteins (2,150) in Exp1.

Sheet 5 (Fig. 1E.Supp): GO enrichment analysis of urea OFF-bead–specific proteins (1,538) in Exp1.

Sheet 6 (Fig. 1F.Supp): GO enrichment analysis shared between urea ON- and OFF-bead proteins (172).

Sheet 7 (Fig. 1G.Supp): Venn diagram protein overlap between urea-on-beads with DIA vs. Sunna.et.al, 2023

Sheet 8 (Fig. 1H.Supp): GO enrichment analysis of proteins (2,528) identified by urea ON-bead DIA

Sheet 9 (Fig. 1I.Supp): GO enrichment analysis of overlapping proteins (1,487) between Sunna et al., 2023 and urea ON-bead datasets.

Sheet 10 (Fig. 1J.Supp): GO enrichment analysis of proteins reported by Sunna et al., 2023

Datasheet 2: Data analysis for Figure 2 Supplementary

Sheet 1 (Supp.Fig.2A): PCA showing distinct clustering by extraction and digestion method of Aldh1l1.

Sheet 2 (Fig. 2B.Supp): Venn diagram comparing proteins identified from SDS lysis ON- and OFF-bead digestion.

Sheet 3 (Fig. 2C.Supp): GO enrichment analysis of proteins (2,896) co-expressed in ON- and OFF-bead fractions.

Sheet 4 (Fig. 2D.Supp): GO enrichment analysis of proteins (933) uniquely identified in SDS OFF-bead digestion.

Sheet 5 (Fig. 2E.Supp): GO enrichment analysis of proteins (262) uniquely identified in SDS ON-bead digestion.

Datasheet 4: Data analysis for Figure 4 Supplementar

Sheet 1 (Fig. 4A.Supp): Volcano plot comparing P2 fraction and homogenate proteomes from Camk2a samples using 8 M urea lysis, in-solution digestion, and DDA LC–MS/MS analysis.
