## Supplementary_Information_Legends for "Deeper neuronal and glial proteomic insights using an optimized pipeline for proximity labeling proteomics"

**FIGURES LEGENDS**
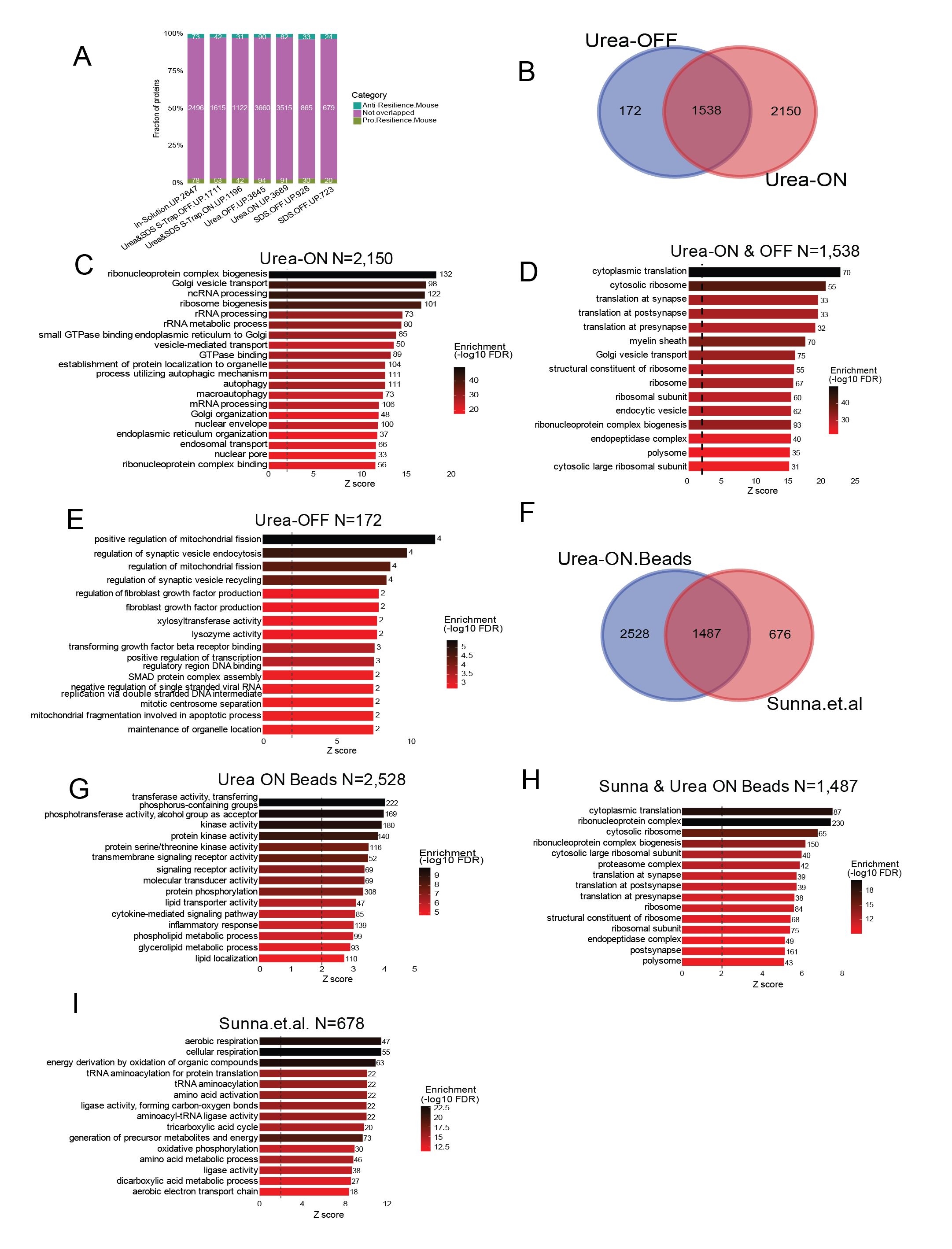
Supplementary Figure 1: **(A) BV2 proteome fractional overlap across resilience:** Stacked bar graph showing the number of overlapped and not-overlapped proteins detected across pro-resilience and anti-resilience different extraction workflows, including Input, ON-beads, OFF-beads. **(B)** **Protein overlaps between Exp1 ON-beads and OFF-beads:** Venn diagram showing the number of shared and unique proteins identified in ON-beads vs OFF-beads fractions. A large, shared core (n=1538) suggests common proteomic coverage, while ON-unique (172) and OFF-unique (2150) proteins indicate preparation-specific enrichment patterns. **(C)** **GO enrichment in Exp4.ON Unique.2150:** Gene Ontology enrichment of proteins uniquely identified in the Exp4 ON-bead dataset (n = 2,150). The ranked bar plot displays the top over-represented biological processes, with bar length indicating Z-score and color intensity representing -log10(FDR) enrichment significance. Highly enriched terms include ribonucleoprotein complex biogenesis, Golgi vesicle transport, ncRNA/rRNA processing, ribosome biogenesis, GTPase-related functions, autophagy, endosomal and nuclear transport pathways. Numbers on the bars denote the count of proteins associated with each GO term. **(D)** **GO enrichment in Exp4.ON.OFF Shared1538:** The bar graph displays top significantly enriched pathways, where Z-score represents enrichment strength and color scale indicates -log10(FDR). Highly represented functions include cytoplasmic translation, cytosolic ribosome, synaptic/post-synaptic translation, myelin sheath organization, Golgi vesicle transport, ribosomal structure, endocytic vesicle pathways, and ribonucleoprotein complex biogenesis. Protein counts associated with each term are shown at the end of the corresponding bars. **(E) GO enrichment in Exp4.OFF.Unique.172:** Gene Ontology enrichment analysis of proteins exclusively detected in the Exp4 OFF-bead fraction (n = 172). The bar plot shows the most significantly enriched biological processes and molecular functions, with bar length representing Z-score and color scale indicating -log10(FDR). Prominent terms include positive regulation of mitochondrial fission, synaptic vesicle endocytosis, regulation of synaptic vesicle recycling, fibroblast growth factor regulation, TGF-β receptor binding, SMAD complex assembly, DNA regulatory region binding, and mitochondrial fragmentation associated with apoptosis. Protein counts for each enrichment term are labeled at the bar ends. **(F)** **Venn for overlapping in Sunna et. al., (year) and Exp4 ON-Beads:** A total of 2,528 proteins were unique to Exp4 ON-beads, 676 were unique to Sunna et al., and 1,487 proteins were shared between both datasets. **(G) GO enrichment in Exp4 ON beads unique 2528**: Enriched pathways for 2,528 ON-bead proteins include kinase transferase activity, signaling, phosphorylation, lipid metabolism, and inflammation **(H) GO enrichment for shared 1487** **proteins in** **Sunna et al and Exp4 On beads**: GO enrichment for the 1,487 proteins shared between Sunna et al. data and Exp4 ON-beads. Top categories include cytoplasmic translation, ribonucleoprotein complex organization, ribosome and proteasome components, and synaptic/post-synaptic translation-related terms. Z-score reflects enrichment of magnitude, and bar color represents -log10(FDR). **(I)** **GO enrichment in 676 were unique to Sunna et al:** Top enriched metabolic and respiratory pathways among the 678 proteins, showing strong enrichment for aerobic or cellular respiration, oxidative phosphorylation, TCA cycle activity, aminoacyl-tRNA ligase functions, and amino acid metabolism. Bar length indicates Z-score; color scale represents -log10(FDR).

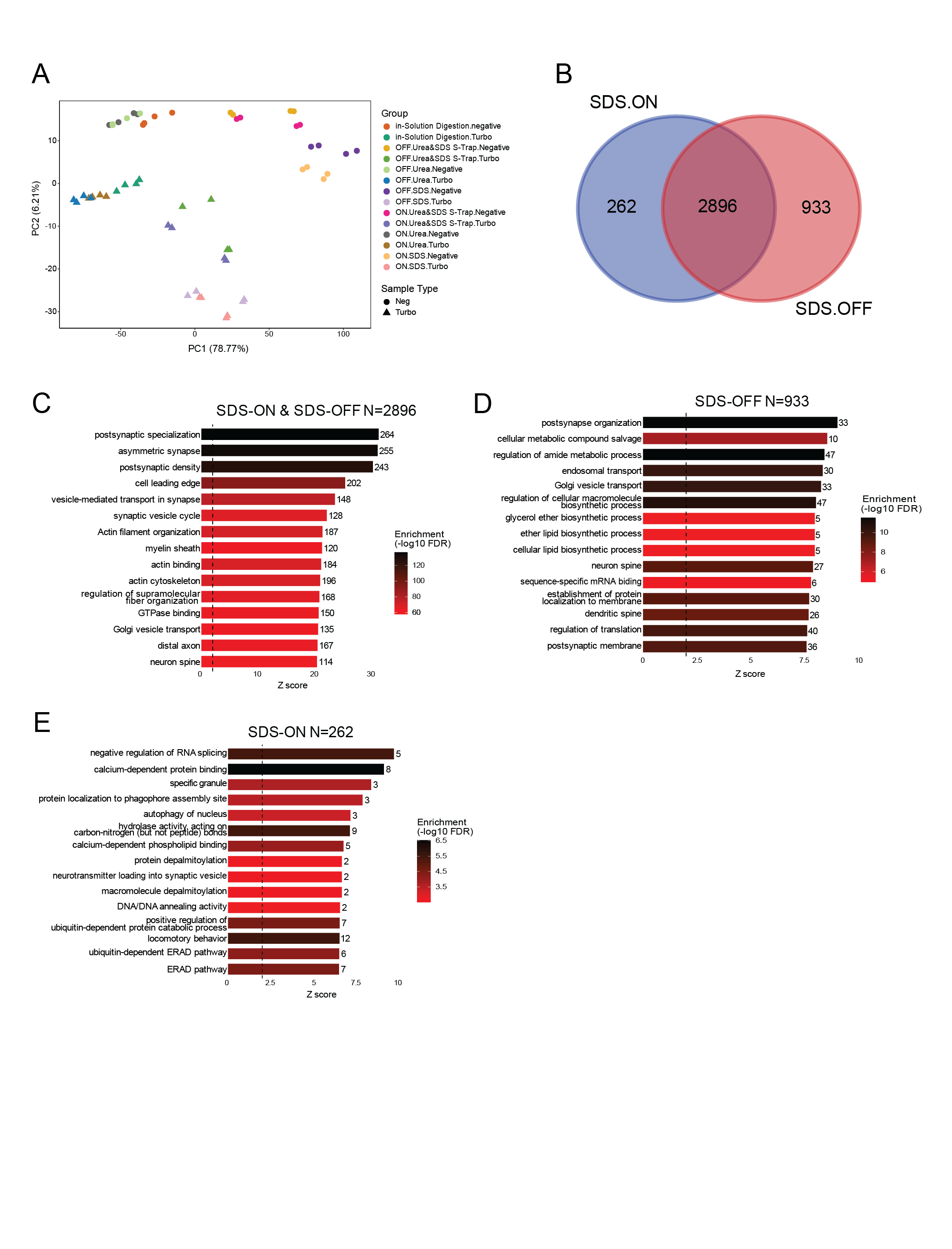

Figure 2 supplementary: **(A) PCA of Aldh1-DDA Samples:** Principal Component Analysis (PCA) of Aldh1-DDA shows clear separation between Turbo and Negative controls across experiments. Biological replicates cluster tightly, indicating high reproducibility and distinct sample type variability. **(B) SDS-ON vs SDS-OFF Protein Overlap:** Venn comparison reveals a large, shared proteome between SDS-ON and SDS-OFF (2,896 shared proteins), with SDS-ON contributing 262 unique proteins and SDS-OFF contributing 933. SDS-OFF yields a broader proteome while maintaining strong overlap with SDS-ON. **(C) GO Enrichment: SDS-ON vs SDS-OFF (2,896 Shared Proteins):** Shared proteins between SDS-ON and SDS-OFF are enriched for neuronal and synaptic pathways, including postsynaptic specialization, vesicle-mediated transport, actin filament organization, synaptic vesicle cycle, and regulation of cytoskeleton dynamics. These functions highlight conserved neuronal signatures independent of digestion mode. **(D) GO Enrichment: SDS-OFF Unique (933 Proteins):** Proteins unique to SDS-OFF digestion are enriched in extracellular matrix organization, cell adhesion, membrane trafficking, and cytoskeletal remodeling processes. SDS-OFF preferentially captures proteins associated with structural and membrane-related functions often missed in SDS-ON. **(E) GO Enrichment: SDS-ON Unique (262 Proteins):** SDS-ON-specific proteins show enrichment in RNA metabolism, translation regulation, ER stress response, and ribonucleoprotein complex assembly. These pathways suggest SDS-ON favors recovery of soluble intracellular and RNA-associated proteins.

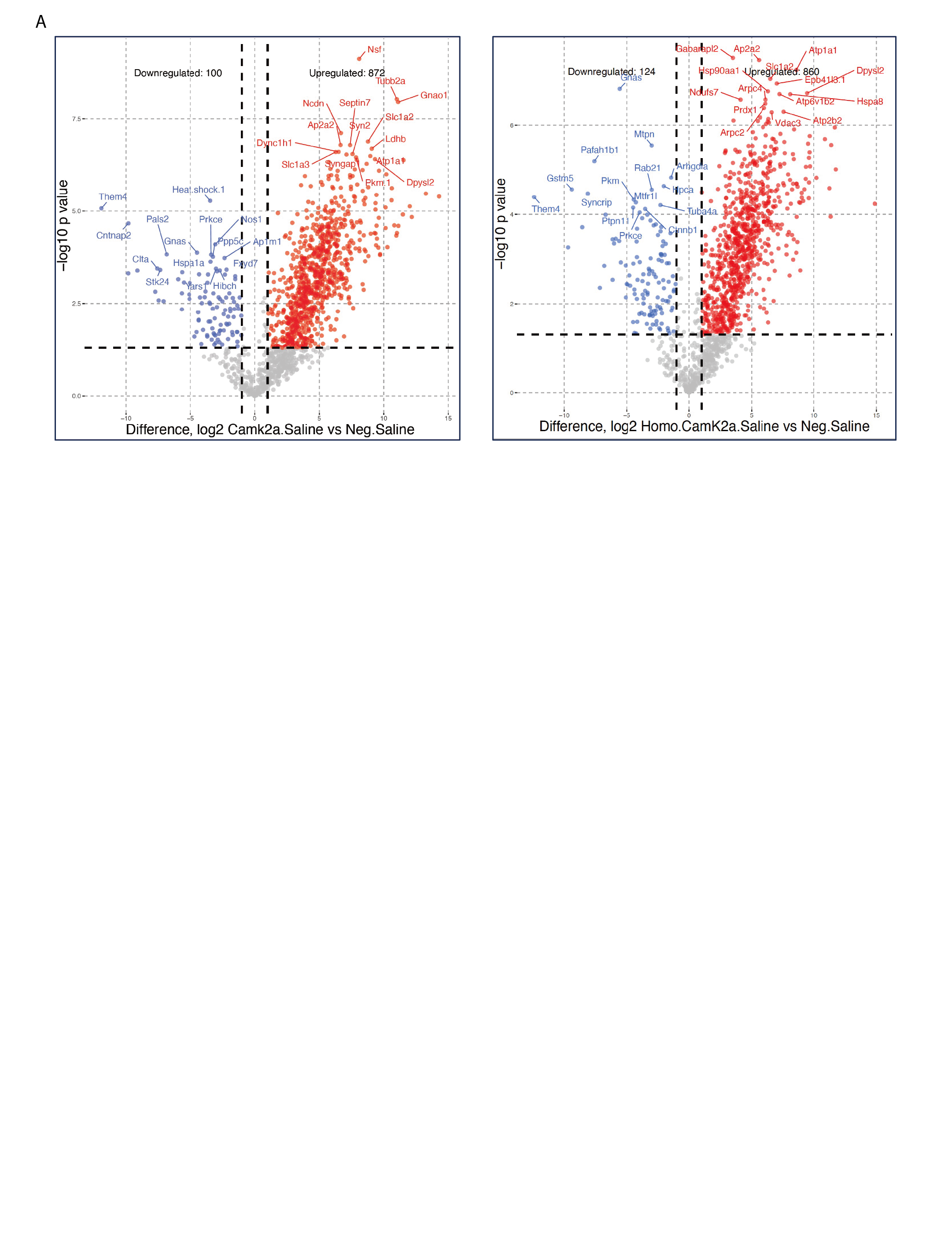

Figure 4 supplementary: **Volcano plots depicting differential protein enrichment in Camk2a-CIBOP samples** relative to Cre-negative controls for whole homogenate (right) and P2 fraction (left). Significantly enriched proteins are highlighted in red (fold change ≥2, p ≤0.05). The optimized DIA workflow identified 860 **proteins** in homogenates and 872 **proteins** in P2 fractions.
